# Inactivation mechanisms of Influenza A virus under pH conditions encountered in aerosol particles as revealed by whole-virus HDX-MS

**DOI:** 10.1101/2022.11.01.514690

**Authors:** Shannon C. David, Oscar Vadas, Irina Glas, Aline Schaub, Beiping Luo, Giovanni D’Angelo, Jonathan Paz Montoya, Nir Bluvshtein, Walter Hugentobler, Liviana K. Klein, Ghislain Motos, Marie Pohl, Kalliopi Violaki, Athanasios Nenes, Ulrich K. Krieger, Silke Stertz, Thomas Peter, Tamar Kohn

## Abstract

Multiple respiratory viruses including Influenza A virus (IAV) can be transmitted via expiratory aerosol particles, and aerosol pH was recently identified as a major factor influencing airborne virus infectivity. For indoor air, small exhaled aerosols undergo rapid acidification to pH ∼4. IAV is known to be sensitive to mildly acidic conditions encountered within host endosomes, however, it is unknown whether the same mechanisms could mediate viral inactivation within the more acidic aerosol micro-environment. Here, we identified that transient exposure to pH 4 caused IAV inactivation by a two-stage process, with an initial sharp decline in infectious titers that was mainly attributed to premature attainment of the post-fusion conformation of viral protein haemagglutinin (HA). Changes to HA were observed by hydrogen-deuterium exchange coupled to mass spectrometry (HDX-MS) as early as 10 seconds post-exposure to acidic conditions. In addition, virion integrity was partially but irreversibly affected by acidic conditions. This was attributed to a progressive unfolding of the internal matrix protein 1 (M1), and aligned with a more gradual decline in viral infectivity with time. In contrast, no acid-mediated changes to the genome or lipid envelope were detected. Our HDX-MS data are in agreement with other more labor-intensive structural analysis techniques such as X-ray crystallography, highlighting the usefulness of whole-virus HDX-MS for multiplexed protein analyses, even within enveloped viruses such as IAV. Improved understanding of respiratory virus fate within exhaled aerosols constitutes a global public health priority, and information gained here could aid development of novel strategies to control the airborne persistence of seasonal and/or pandemic influenza in the future.

## INTRODUCTION

Airborne transmission of infectious agents relies on the ability of pathogens to survive aerosol transport whilst they transit between hosts. Evidence in the literature increasingly points towards the expiratory aerosol as a major transmission mode for many respiratory viruses, including SARS-CoV-2 (1–7), human rhinovirus (8–11), respiratory syncytial virus (12), human adenovirus (13), and influenza A virus (IAV) (14–23). The aerosol route dominates under certain environmental conditions, particularly indoor environments that are poorly ventilated. In one study, airborne transmission was estimated to account for half of all IAV transmission events within households (19). Viruses are also vulnerable during airborne transit, however, there continues to be a fundamental lack of understanding as to mechanisms that could drive virus inactivation within expiratory aerosols.

Upon exhalation, aerosols rapidly lose moisture and heat, with a large concomitant change in volume as equilibrium is established with the indoor environment. Numerous studies have attempted to assess how this process affects viral infectivity, with temperature, salt concentration, relative humidity, presence of respiratory surfactants, and air/liquid interface interactions all postulated to play a role (24–27). Importantly, aerosol pH has recently been recognized as a principle determinant of virus fate within aerosols (28,29). When exhaled into typical indoor air, the pH of exhaled aerosol particles quickly deviates from neutral, and decreases to ∼4 (29,30). Such a low pH can trigger the inactivation of acid-sensitive viruses. IAV in particular is sensitive to acidic conditions encountered in the endosome (37°C, ∼pH 5.5), with pH-induced conformational change to HA driving this sensitivity (31,32). The pH-activated form of HA – also referred to as post-fusion conformation – allows HA to mediate membrane fusion between the endosome and the viral envelope, and allows viral uncoating and entry to proceed. However, if the post-fusion conformation is attained prematurely, HA will be unable to bind host cells and the virus is effectively inactivated. It is yet to be confirmed whether the same mechanism drives IAV inactivation under the more acidic conditions found within exhaled aerosols. Aerosol pH is within the range of general protein unfolding induced by acids (occurs between pH 2 and 5, whilst base-induced protein unfolding usually requires pH 10 or higher (33)), and whether IAV inactivation within aerosols could be due to global protein denaturation or specifically due to HA remains to be understood.

In this study, we investigated the effects of aerosol-associated pH on IAV structural integrity, and determined the resulting effects on the viral infectious cycle. To decouple pH effects from other factors present within an evaporating aerosol micro-environment, we performed this study in bulk solutions with controlled pH values representative of aerosol particles in indoor air. IAV also presents certain challenges for structural analysis, as it is a complex virion comprised of 11 individual proteins, some of which are membrane embedded. To gain high-resolution information on potential changes occurring to IAV proteins under acidic aerosol conditions, we utilized hydrogen-deuterium exchange coupled to mass-spectrometry (HDX-MS). This method assesses local as well as global conformational changes of proteins in their near-native environment. By incubating a protein of interest in deuterated buffer (D_2_O), amide hydrogens will passively exchange for deuterium, and the rate of exchange is dependent on the structural environment of individual amides (34). Changes in local protein structure can therefore be distinguished by comparing amide exchanges rates of equivalent peptides after exposure of proteins to different conditions (35). Importantly, this technique allows investigation of proteins in the whole-virion conformation (36–41), as deuterium will diffuse as much as normal water through an intact viral capsid.

Two prior studies have applied HDX-MS to whole IAV, though both focused on the effect of host endosomal pH (∼5.5) on the surface-exposed HA protein (42,43). We sought to multiplex our HDX-MS analysis by analyzing three different viral proteins at once; surface-exposed HA, the mid-layer matrix protein 1 (M1), and the internal nucleoprotein (NP). Internal IAV proteins have never been characterized using HDX-MS before, under native nor acidic conditions. To complement the study on protein structure and obtain a global analysis of the state of IAV, integrity of the viral genome and of viral lipids were also assessed.

## RESULTS

### IAV is rapidly inactivated under expiratory aerosol conditions

The majority of exhaled aerosols from humans are <5 μm, with a large fraction being <1 μm for most respiratory activities (8,44). Previous studies have specifically identified influenza virus in aerosols of these sizes from infected individuals (23,45,46). Upon exhalation, aerosols rapidly lose water and equilibrate with acidic and basic trace gases present in indoor air, including nitric acid and ammonia. We previously published a biophysical Respiratory Aerosol Model (ResAM) used to show an exhaled particle of 1 μm initial radius will acidify to pH ∼4 due to this equilibration (29). Depending on the initial size of the particle, the pH development will vary. Here, ResAM was used to estimate the pH trajectory of exhaled particles of 3 different initial sizes (0.2 μm, 1 μm, and 5 μm radius), representative of the range of ‘fine’ aerosols produced by humans. ResAM predicted that the same terminal acidic pH of ∼4 would be reached for all 3 particle sizes, with the smallest modelled particle size reaching this point less than 10 seconds post-exhalation (**Fig. 1A**). With larger particles, more time is needed for the diffusion of trace acids into the particle, thus the acidification process was predicted to be slower, though a terminal pH of ∼4 was reached in all cases.

**Figure 1.**
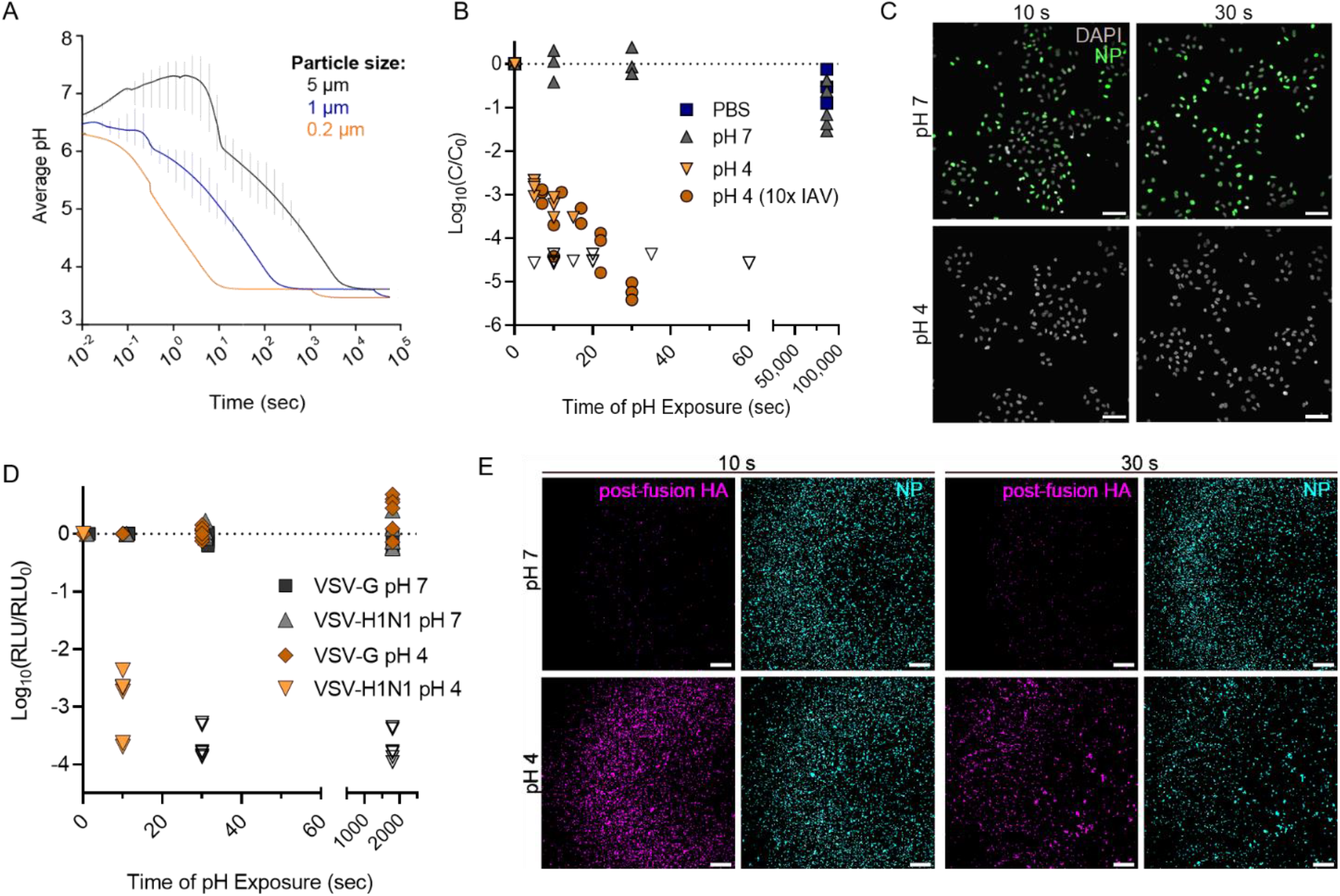
Acidification of an exhaled aerosol, and the effect on IAV infectivity. **A)** Average pH within an exhaled respiratory aerosol particle as it transitions from the nasal microenvironment to typical indoor air, modelled using ResAM. The particle evaporates with time (in seconds), resulting in acidification down to ∼pH 3.8. Progressive changes in pH are modelled for particles of 3 different initial sizes. Vertical error bars indicate the pH range between the innermost and outermost aerosol shells at each timepoint, with the mean pH presented as a continuous line. Thermodynamic and kinetic properties are those of synthetic lung fluid (SLF), a representative respiratory matrix. The indoor air conditions were set at 20°C and 50% RH, and the exhaled air is assumed to mix into the indoor air using a turbulent eddy diffusion coefficient of 50 cm^2^ /s. **B)** Inactivation of IAV due to exposure to acidic pH 4. IAV was spiked into neutral solutions of PBS or pH 7 buffer, or spiked into acidic pH 4 buffer. Aliquots were taken at multiple time-points post spiking (up to 86,400 sec, or 24 hours), and neutralized for quantification of remaining viral titer by plaque assay. For pH 4 samples, virus was spiked at 10^6^ PFU/mL, or at 10^7^ PFU/mL (10x IAV), to demonstrate that viral concentration did not affect the rate of viral decay. Data points indicate the mean ± SD of triplicate measurements at each time-point, presented as log_10_ of remaining concentration (C) at each time-point relative to the initial concentration (C_0_). Data is compiled from 3 independent experiments, data points where C was below the limit of detection for each plaque assay are indicated by open symbols. Dotted line indicates no decay. **C)** Immunofluorescence images of A549 cells infected with IAV samples treated at pH 4 or 7 for 10 or 30 seconds. All samples were neutralized prior to cell infections. A549 cells were infected at a multiplicity of infection (MOI) of 2, and IAV protein production was visualized by immunofluorescent staining of nucleoprotein (NP) at 3 hours post-infection. Scale bar corresponds to 75 μm. Images representative of duplicate experiments. **D)** VSV particles encoding Renilla Luciferase were pseudotyped with HA and NA of the A/WSN/33 virus (VSV-H1N1), for comparison to VSV expressing the wild-type surface protein G (VSV-G). Both VSV stocks were acidified to pH 4, and samples were neutralized after 10 or 30 sec of pH exposure. Control samples of each VSV strain were held at pH 7. Entry capacity of treated VSV samples was then determined by measuring the luciferase activity 7 hours post infection in MDCK cells. Data was derived from 3 independent experiments, with samples quantified in duplicate. Data is presented as log_10_ of relative light units (RLU) at each time point relative to RLU of non-treated samples (RLU_0_). Dotted line indicates no decay. **E)** IAV particles were pH-treated and neutralized as in C, then stained with an HA-specific antibody recognizing the post-fusion HA conformation (purple). Staining with an NP-specific antibody (blue) served as a control to ensure the same amount of viral material was deposited between samples. Scale bar corresponds to 250 μm. Images representative of duplicate experiments.

We first investigated the kinetics of IAV inactivation under conditions representative of aerosols equilibrated with indoor air (room-temperature, pH 4). Infectivity data (**Fig. 1B**) show that transient exposure to pH 4 at room temperature results in rapid virus inactivation, with infectious viral titers decaying by approximately 3-log_10_ within 10 seconds of pH 4 exposure, and by >5-log_10_ by 30 seconds. Conversely, IAV spiked into pH 7 solution or PBS (both neutral controls) and held at the same temperature showed minimal decay for the full 24-hour monitoring period. Virus samples exposed to pH 4 also showed no productive infection in host epithelial cells at 3 hours (**Fig. 1C**) or 6 hours post-infection (**Supp. Fig. S1**) when compared to neutral controls, suggesting that an early step of the viral infection cycle was specifically impaired by low pH treatment.

To investigate whether this viral decay could be mediated specifically by HA (rather than general protein unfolding), pseudotyped Vesicular stomatitis virus (VSV) strains were used. VSV was pseudotyped with H1N1 IAV surface proteins HA and neuraminidase (NA), generating VSV-H1N1. Inactivation kinetics of the pseudotyped VSV strain at pH 4 were very similar to inactivation of wild-type IAV, with ∼3-log_10_ decrease in infectivity after 10 seconds pH-exposure, and residual infectivity being below detection limits by 30 seconds (**Fig. 1D**). As IAV NA does not function in viral attachment or uncoating inside host cells, this result indicates that the majority of virus inactivation for pseudotyped VSV was mediated by non-reversible changes to HA. Control VSV, which expresses wild-type G protein rather than HA for host-cell attachment, showed no loss of infectivity as a result of the same pH treatment. Importantly, VSV G protein undergoes a similar low pH conformational change like IAV HA to mediate envelope fusion with host cells, but the change to G protein is entirely reversible (47). Additionally, fluorescence imaging of whole pH-treated IAV particles showed increased staining for the post-fusion form of HA after 10 seconds of pH 4 exposure (**Fig. 1E**), whilst the post-fusion form of HA was minimally present on pH 7-treated viruses. Some low-level signal was observed at pH 7, however this was attributed to background staining.

### Transient acidic exposure induces localized conformational changes in external IAV protein haemagglutinin (HA)

Data in **Figure 1** suggested the loss of IAV infectivity in the pH 4 condition was indeed due to a conformation change of HA into its post-fusion form, rather than general protein denaturation. To confirm this and map specific regions of HA affected by pH 4 exposure, we utilized whole-virus HDX-MS. Intact IAV particles were rapidly acidified to pH 4 in bulk solution prior to neutralization, whilst controls were held in neutral conditions for the same exposure time. All samples were then incubated in D_2_O buffer on ice for a dedicated time before H/D exchange reactions were quenched to lock exchanged deuterium in place. During quenching, whole viruses were also lysed with a urea and formic acid mixture, releasing internal proteins for subsequent protease digestion and HDX-MS analysis. Viral lysis was incorporated as the final step to retain the virus in its near native conformation during pH treatment, neutralization, and D_2_O incubation (HDX-MS schematic in **Fig. 2A**).

**Figure 2.**
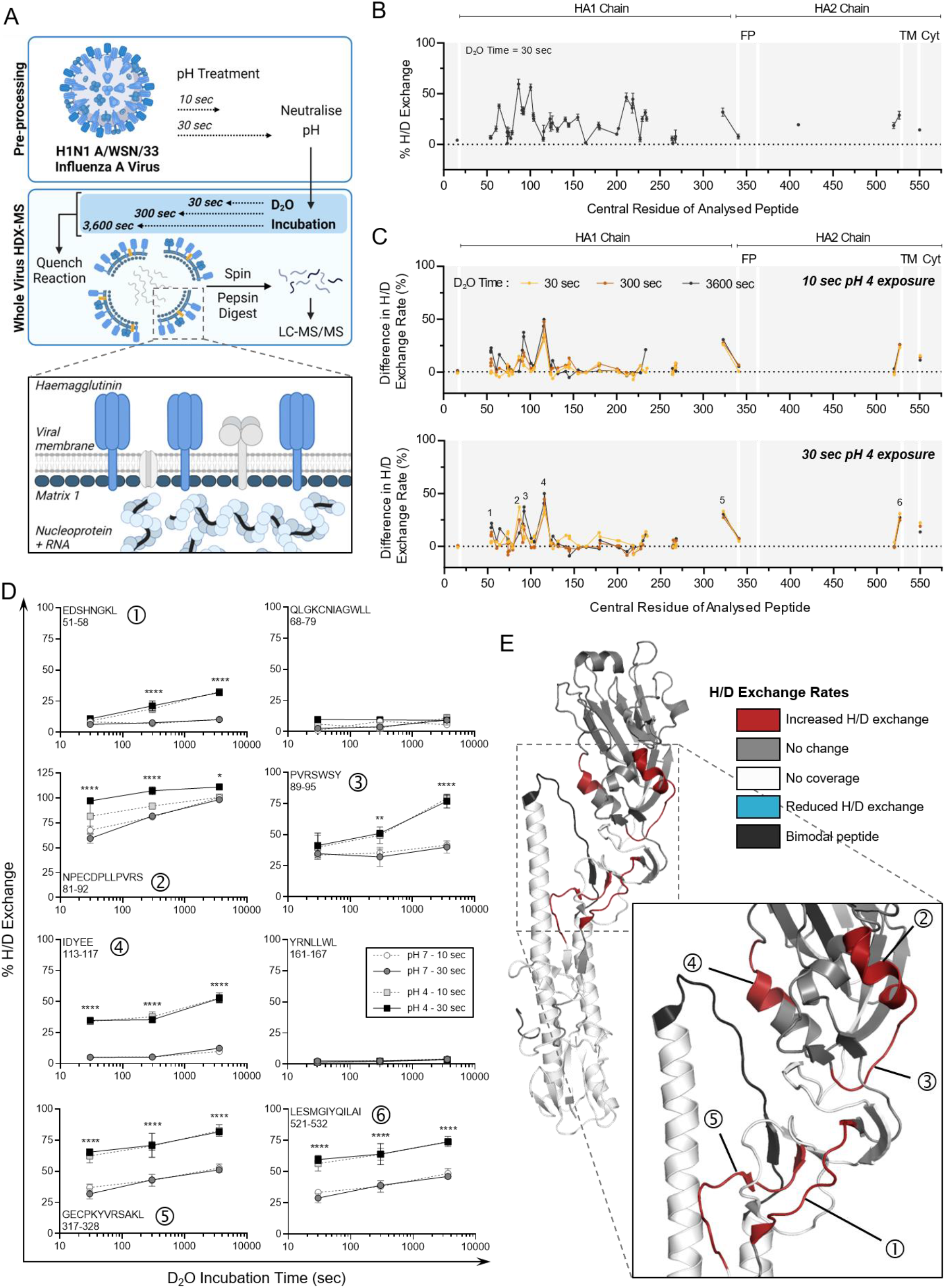
HDX-MS characterization of HA protein from whole IAV after transient exposure to acidic conditions. **A)** Schematic of the whole-virus HDX-MS process. In bulk solution, whole virus is held at neutral pH or is acidified at pH 4 for 10 or 30 seconds prior to neutralizing back to pH 7. Neutralized whole-virus samples are incubated in D_2_O buffer for 30, 300, or 3,600 seconds, then quenched, prior to pepsin digestion and LC-MS analysis. Organization of viral proteins within the intact influenza A virion are highlighted by the inset, with the external Haemagglutinin (HA), and the internal proteins Matrix 1 (M1) and Nucleoprotein (NP) colored blue for visibility. **B)** Hydrogen/deuterium (H/D) exchange percentages measured for HA protein from whole IAV samples held at neutral conditions (pH 7) for 30 seconds. Samples were then incubated in D_2_O for 30 seconds, quenched, and digested prior to HDX-MS analysis. Data points indicate the mean percentage of H/D exchange ± SD for triplicate samples. Peptides are organized linearly according to the position of the central amino acid of each individual peptide along the published sequence for HA (Uniprot: P03452). Continuous lines indicate regions of continuous peptide coverage by MS. Peptides showing bimodal activity are excluded from the graph. Protein domains of interest are indicated by grey shading and white lines; FP = fusion peptide, TM = transmembrane domain, Cyt = cytoplasmic region. **C)** Residual plots showing the difference in H/D exchange percentages induced by exposure to pH 4 for 10 seconds or 30 seconds, when compared to corresponding pH 7 control samples. Individual data points indicate the mean percentage of H/D exchange difference for triplicate samples. Dotted line indicates no difference in H/D exchange. Peptides of interest showing increased H/D exchange are numbered 1 – 6. **D)** Uptake plots for selected Peptides 1-6 showing increased H/D exchange after pH 4 exposure. Data shows the total percentage of H/D exchange for pH 7 or 4 samples after D_2_O incubation for 30, 300, or 3,600 seconds. The sequence and residue positions of each peptide are indicated in each panel. Data points indicate the mean percentage of H/D exchange ± SD for triplicate samples. Data analyzed by Two-way ANOVA (** p < 0.01, *** p < 0.001, **** p < 0.0001, when comparing 30 second pH-exposed samples for each D_2_O time-point. Statistical results for comparison of 10 second pH-exposed samples appear in **Supplementary Table S1**). **E)** Protein regions showing increased (red) or decreased (blue) H/D exchange rates, or showing no change in H/D exchange rates (grey) after 30 seconds at pH 4 (compared to pH 7 control) mapped onto the crystal structure of A/PR8 HA (PDB: 1RU7). White indicates regions of no peptide coverage, and peptides with bimodal activity are colored black. HA monomer is shown, with inset showing the specific location of Peptides 1-5, corresponding to those in panel D. Peptide 6 (521-532) from panel D is not indicated as the crystal structure is truncated prior to these residues.

The HA protein itself is present as a homotrimer on the surface of whole IAV, with a membrane-embedded region and internal region anchoring the protein to the viral matrix layer (**Fig 2A**). Each HA monomer is comprised of two subunits, HA1 and HA2, connected by a short and relatively flexible hinge region. IAV in the neutral pH 7 condition was first used to characterize the global D_2_O incorporation levels of this surface-exposed glycoprotein. HA showed an alternance of fast- and slow-exchanging regions, suggesting the presence of dynamic and more structured regions (**Fig. 2B**). Peptide coverage was high for the HA1 region and much reduced for membrane-associated HA2 region. Reduced coverage for HA2 has been observed in prior HDX-MS studies analyzing HA of a different IAV strain (42,43). Overall, 74% of residues within HA1 and 30% of those within HA2 of IAV were covered by our whole-virus HDX-MS analysis (**Supp. Fig. S2**). Peptides analyzed in our study showed good correlation in their deuteration levels with previous study of HA by HDX-MS (42), with the same alternance in high- and low-exchanging regions despite use of different virus subtypes and strains (H3N2 X-31 vs H1N1 A/WSN/33).

Following characterization of HA dynamics at neutral pH, H/D exchange rates after transient pH 4 treatment of the whole virus were characterized. Three different D_2_O incubation times (30 sec, 300 sec, and 3,600 sec) were used here to allow detection of pH-induced changes occurring in highly flexible as well as in more structured regions. **Figure 2C** shows comparison of H/D exchange rates between neutral and pH 4 conditions, which identified several peptides of interest displaying increased exchange post-acid treatment (see **Supp. Table S3** for exchange levels in all detected HA peptides). These peptides with increased exchange were detected after just 10 seconds at low pH, and increasing the pH exposure time to 30 seconds did not appear to amplify these differences. Neutralization of all samples prior to HDX-MS ensured these pH-induced changes were non-reversible, and thus are likely to indicate the state of the HA protein on whole IAV after re-inhalation and neutralization of an acidified aerosol on the mucus membrane of a new host.

Six discrete peptides showed significant differences as a result of pH, 5 of which were localized within the HA1 chain (peptides numbered in bottom panel of **Fig. 2C**). The first identified peptide was from residue 51-58 (Peptide 1 in **Fig. 2D**). H/D exchange rates for this peptide after 30 seconds of D_2_O incubation were comparable between pH conditions (pH 4 vs 7), though significant differences emerged after longer D_2_O exposures of 300 and 3,600 seconds (see **Supp. Table S1** for all statistical comparisons). Peptide 1 is located within the fusion domain (F’) of the HA1 subunit. Elevated H/D exchange here after pH 4 treatment is likely due to disruption of stabilizing contacts between this domain and the proximally located HA2 subunit [reported in (48)] as the protein transitions to a post-fusion conformation. Peptide 5 (residues 317-328) is located in the same fusion region, and showed a similar enhancement of H/D exchange post-pH 4 treatment. Two peptides that were unaffected by pH are also included in **Fig. 2D**, to highlight that H/D exchange rates were altered in specific parts of the HA protein only.

Peptides 2 (residues 81-92) and 3 (residues 89-95) are overlapping, though showed a differential kinetic in detected H/D exchange, suggesting that the 80-90 region is more dynamic than the 90-95 region. These observations were not anticipated as the high-resolution X-ray crystal structure of HA [PDB: 1RU7 from (49)] suggests the presence of an α-helix in the 80-90 region, which was expected to be more protected from H/D exchange than the unstructured 90-95 region (positions highlighted in **Fig. 2E**). Peptide 4 (residues 113-117) lies in the vestigial esterase (E′) subdomain of HA1 and spans an alpha-helical region and loops. Highly significant increases in H/D exchange rate for this peptide were observed at every D_2_O incubation time, suggesting that the secondary structure element might rapidly unfold upon pH 4 treatment. Peptide 4 is also buried within the center of the HA homotrimer in the pre-fusion conformation, making close contact with the β-loop of the HA2 subunit in neutral conditions. In the post-fusion confirmation, the RBD of HA1 is rotated away from the central stalk (43,50), releasing contact with HA2 and leaving these previously protected peptides more exposed to the external environment.

Peptide 6 (residues 521-532) sits immediately prior to the anchoring transmembrane region of HA2. During transition to post-fusion HA, this membrane proximal region is reported to become disordered (50), which is in-line with the large increase in H/D exchange rate observed after pH 4 exposure here. A similar increase in H/D exchange within the C-terminal region of HA for the H3N2 IAV strain X-31 was observed post-fusion by Garcia *et al*. (42). Interestingly, we observed one peptide within HA showing bimodal distribution, also referred as EX1 kinetics (residue 402-418, colored black in **Fig. 2E**), which is addressed in the next section.

### Peptide bimodal distribution is induced by acidic exposure for the β-loop of HA protein

One peptide (HA2, 402-418) showing bimodal H/D exchange behavior was detected after pH 7 and pH 4 treatments, suggesting the presence of distinct populations with different conformations. This peptide is located within the β-loop of the HA2 subunit (**Fig. 3A**), which is one of the primary regions known to modify its conformation in pH-triggered HA, changing from a flexible loop to a stabilized α-helix. When in neutral conditions, unimodal H/D exchange was initially detected for this peptide. Representative spectra (**Fig. 3B**) show the mass shift and fraction of each population for the 402-418 peptide after a 30 second D_2_O incubation, with a single peak indicating unimodal distribution for pH 7 samples. A second population (orange) then became distinguishable after longer D_2_O incubation times, indicating bimodal distribution, though these two peaks were mostly overlapping for the entire D_2_O incubation period (up to 1 hour). Conversely, this bimodal distribution was observed even after short D_2_O incubation times following pH 4 treatment of the whole virus. The second peak was also substantially more deuterated than in control samples. Deconvolution analysis was applied to quantify the fraction of each population, and showed that this second, highly deuterated population dominated after exposure to pH 4 conditions (**Fig. 3C**).

**Figure 3.**
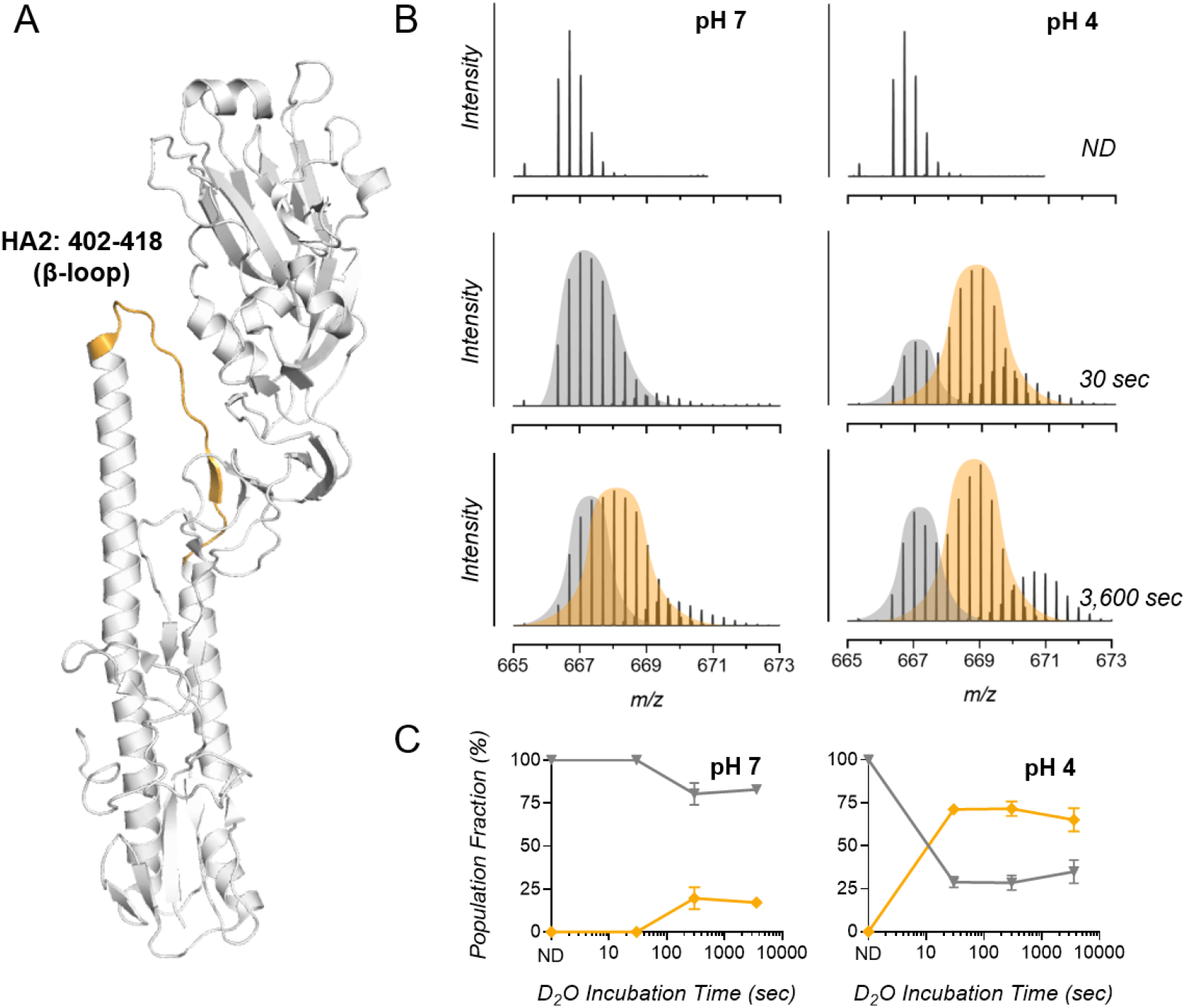
Regions of bimodal H/D exchange within IAV HA identified by whole virus HDX-MS. **A)** HDX-MS analysis revealed the HA peptide 402 – 418 (part of the HA2 subunit) displayed bimodal activity, and this peptide is highlighted orange on the crystal structure of A/PR8 HA (PDB: 1RU7). **B)** Representative spectra of triplicate samples from pH 7 and pH 4 groups showing a mass shift for bimodal peptide 402 – 418 with increasing D_2_O incubation times. Mass shift is compared to samples incubated in non-deuterated buffer (ND). Bimodal envelopes are separated by grey and orange coloring. **C)** Proportions of the left (grey) and right (orange) populations are graphed with time. Data indicates the mean ± SD for triplicate samples analyzed by HDX-MS.

Bimodal exchange behavior is indicative of the presence of two distinct conformations, with long-range allosteric rearrangements occurring upon transiting from one state to another. Benhaim *et al*. (43) reported a similar observation for the HA β-loop of the H3N2 strain X-31 after acidification at pH 5 at a host-temperature of 37°C. Their study showed appearance of an intermediate and highly deuterated peak after ∼1 minute of pH exposure, which progressively disappeared to be overtaken by a highly protected segment as pH exposure progressed. In our conditions, exposure times are too short to show this progressive disappearance of the second peak, and indicates HA is in the very initial stages of conformational change.

### Transient exposure to low pH has a small effect on virion integrity, but not on genome integrity or lipid content

In addition to monitoring conformational changes on the surface-exposed HA trimeric protein, we determined whether structural integrity of virus particles and of genomic material was affected when whole IAV was pH-exposed. We suspected additional mechanisms of inactivation were occurring as HA conformational changes observed by HDX-MS were very similar upon 10- or 30-seconds of pH 4 exposure, though infectious titers continued to drop (**Fig. 1B**).

Initially, the structure of the viral capsid was assessed post-pH treatment by TEM imaging, and showed that IAV in neutral conditions (PBS and pH 7) was spherical/ovoid, with virions relatively well dispersed across the grids (**Fig. 4A**). Spikes of HA were also readily visible and well organized across the viral surface. In contrast, after exposing the virus to pH 4 (30 seconds), virions were aggregated, and those appearing as single particles or smaller clusters had reduced structural integrity. Most noticeably, the organization of HA spikes on the surface of individual virions was compromised after pH treatment, providing additional evidence that HA had undergone an irreversible conformational change following transient acidic exposure. The post-fusion HA trimeric complex is known to expose hydrophobic regions within the HA2 stem that are buried pre-fusion (51–53). Increased hydrophobicity of virions due to HA conformation change is likely to mediate the partial aggregation of viral particles observed at pH 4. It is important to note our prior work has established that viral aggregation is not the driver of infectious titer loss in our system (29). Some virions also appeared to be in the process of lysis by TEM, though overall, most viral particles appeared structurally intact after pH 4 treatment.

**Figure 4.**
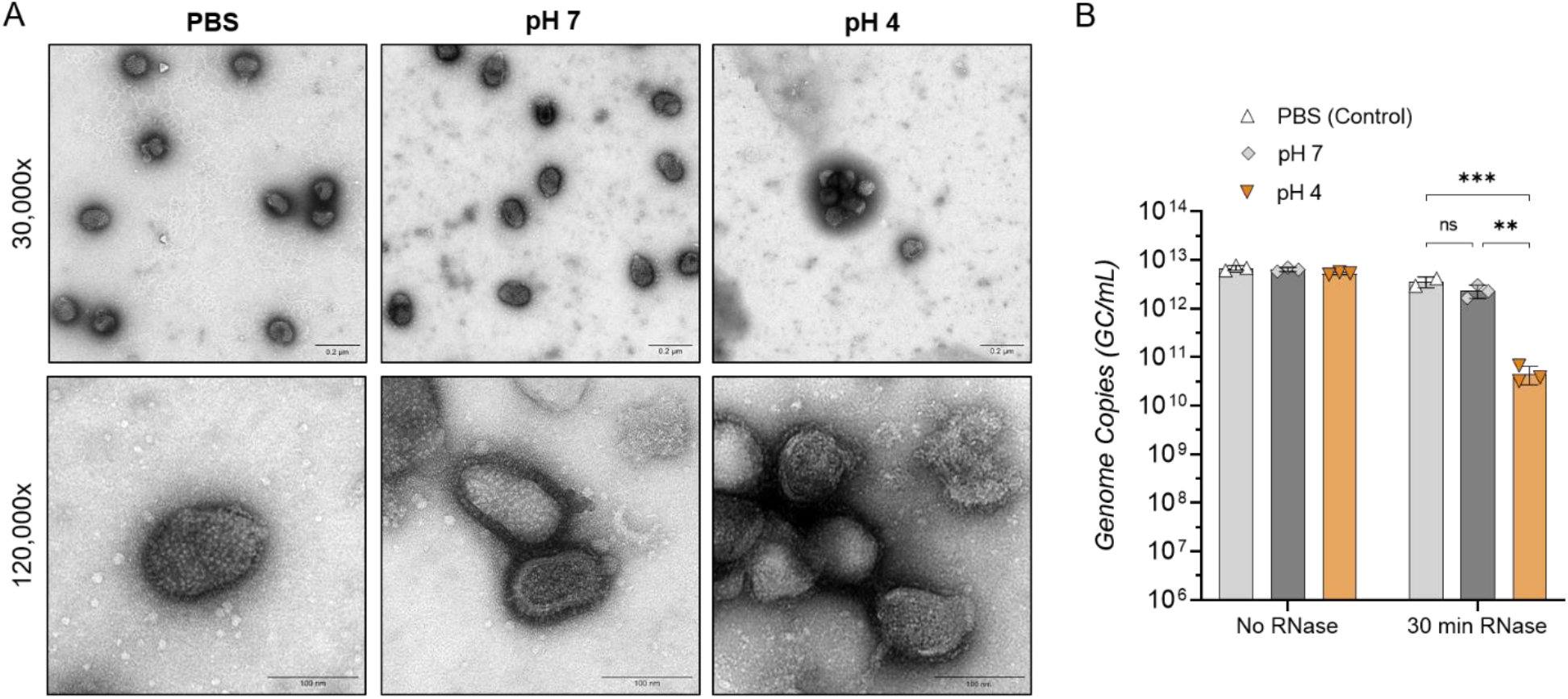
Effect of aerosol pH on IAV structural integrity. IAV was adjusted to pH 4, then neutralized back to pH 7 after 30 seconds, control samples were mock-adjusted to pH 7 for the same time period, or were suspended in PBS only. **A)** Neutralized and control samples were fixed in PFA, then stained with 2% uranyl acetate and imaged using the Tecnai Spirit TEM (imaged at 30,000x and 120,000x magnification). Images are representative of duplicate samples. **B)** Alternatively, neutralized virus samples were incubated in the presence of RNase or PBS (no RNase control) prior to RNA extraction and genome quantification by RT-qPCR. Data presented as mean genome copies/mL ± SD for triplicate samples. Data analyzed by Two-Way ANOVA (** p < 0.01, *** p < 0.001, ns not significant).

Virion integrity was additionally investigated by determining if RNA release occurred during pH treatment. Whole viral samples were exposed to pH 4 or pH 7 for 30 seconds, prior to neutralization back to pH 7. Samples were then split in 2 groups; one treated with RNase, and the second mixed with PBS (no RNase) before incubation of all samples at 37°C for 30 min. Samples were then subject to viral RNA extraction and quantification by RT-qPCR. Samples with no RNase treatment showed comparable genome amplification regardless of pH pre-treatment (**Fig. 4B**), indicating the segment of the IAV genome amplified here was not affected by pH alone. Additionally, exposure of whole virus to acidic pH for up to 30 minutes resulted in no change to the amplified genome segment (**Supp. Fig. S3**). For samples mixed with RNase however, we observed varying levels of amplifiable RNA copies. Copies in PBS and pH 7 control samples did not drop significantly after RNase treatment, indicating the amplified genome segment was sufficiently protected from degradation by a whole, intact virion. Conversely, pH 4-treated samples showed a significant loss of amplifiable genome after RNase exposure, suggesting some virions were no longer intact to protect the genome from RNase-mediated degradation. Overall, the drop in total genome copies was approximately 2-log_10_ for pH 4-treated samples after 30 seconds. Whilst significant, this is not the same magnitude of titer decrease observed for infectivity (upwards of 5-log_10_ loss of viral titer in the same time-frame), indicating that loss of virus integrity is likely to be contributing to inactivation, but is not the primary driver.

Finally, we assessed if pH treatment could affect the lipid classes within the IAV lipid envelope. Here, we measured the relative abundance of 500 lipid species using liquid chromatography/mass spectrometry (LC/MS) untargeted lipidomics, after whole-virus exposure to pH 4, 5, or 7 for 30 seconds. Data in **Supplementary Figure S4** shows no significant change in lipid levels upon pH 4 or pH 5 treatment over the time-frame of interest. While key viral lipids (e.g. sterols) have not been considered here, and future work will assess potential lipid alterations in more detail, data indicates there are no major alterations to the lipid envelope composition after acidic pH exposure over the time-scale of interest.

### Ambient acidic exposure has no effect on internal nucleoprotein (NP)

As some virions were ruptured as a result of pH treatment, it was of interest to determine whether the conformation of the internal protein NP was also affected. Here, the multiplex capabilities of whole-virus HDX-MS were useful, as the same set of samples used for prior HA analysis were now analyzed for structural changes to NP. At neutral conditions (pH 7), NP showed a number of inherently flexible regions, particularly towards the C-terminal end of the protein (**Fig. 5A**). When comparing pH 7 to pH 4 treated samples, no significant changes in H/D exchange rates were detected for any peptides within NP (**Fig. 5B – C**, and **Supp. Table S4** for full list of NP peptides), indicating no conformational changes or localized unfolding were occurring despite partial viral lysis and exposure to acidic conditions. NP gave the highest peptide coverage of this study, with 75% of the entire sequence covered by at least one peptide (**Supp. Fig. S5**). It cannot be excluded that non-covered regions may show pH-induced conformational changes, and this could be the focus of follow-up work.

**Figure 5.**
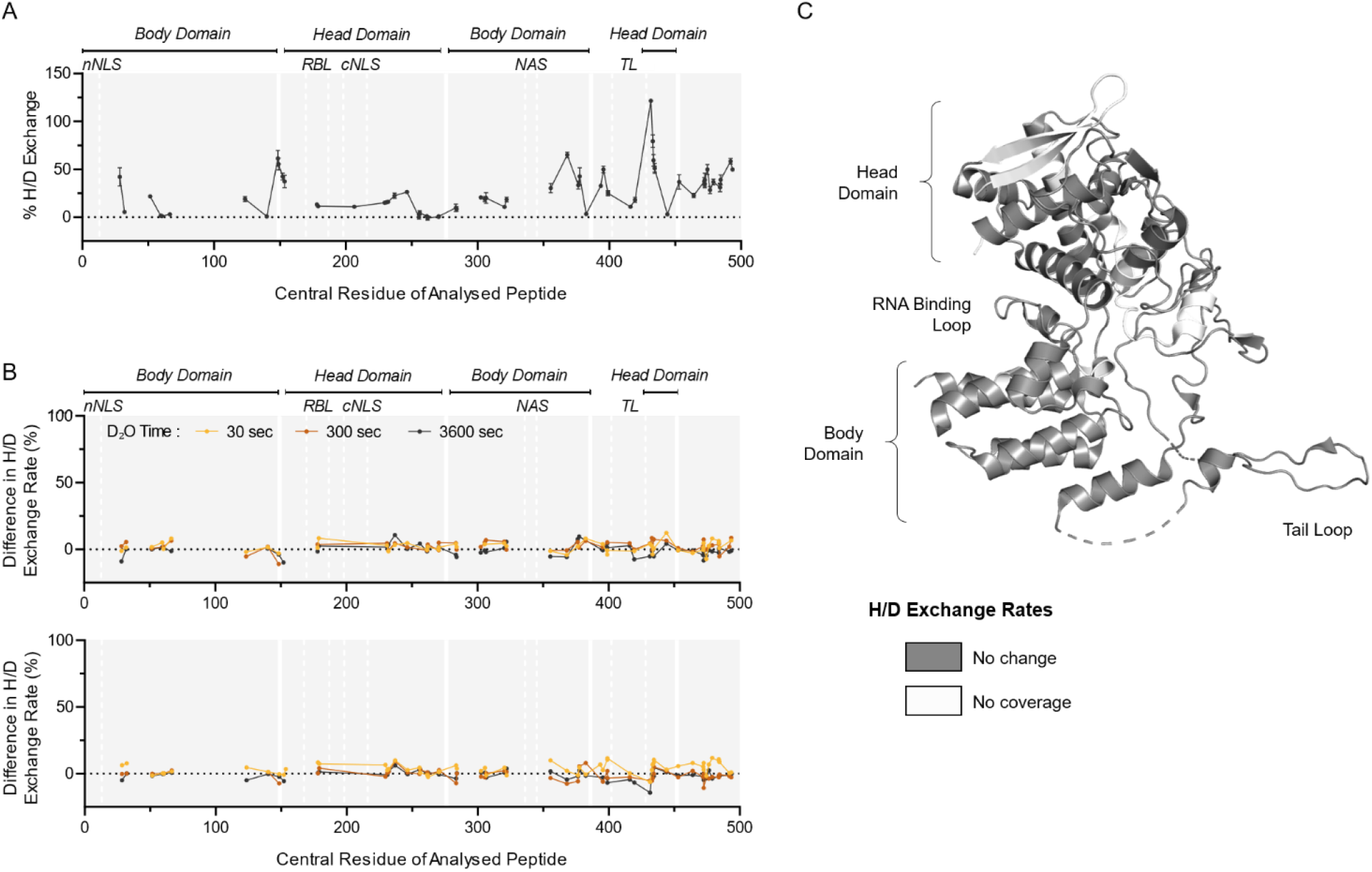
HDX-MS characterization of NP protein from whole IAV after transient exposure to acidic conditions. **A)** H/D exchange percentages measured for NP protein from whole IAV samples held at neutral conditions (pH 7) for 30 seconds. Samples were then incubated in D_2_O for 30 seconds, quenched, and digested prior to HDX-MS analysis. Data points indicate the mean percentage of H/D exchange ± SD for each peptide plotted using its central amino acid (n=3, Uniprot: P03466). Continuous lines indicate regions of continuous peptide coverage by MS. Protein domains of interest are indicated by grey shading and white lines; nNLS = non-conventional nuclear localization signal, RBL = RNA binding loop, cNLS = conventional nuclear localization signal, NAS = nuclear accumulation signal, TL = tail loop. **B)** Whole virus samples were held at neutral pH, or adjusted to pH 4, then neutralized back to pH 7 after 10 or 30 seconds prior to processing for HDX-MS. Data presented as residual plots showing the difference in H/D exchange percentages induced by exposure to pH 4 when compared to corresponding pH 7 control samples. Individual data points indicate the mean percentage of H/D exchange difference for triplicate samples. **C)** Crystal structure of A/WSN/33 NP (PDB: 2IQH), showing regions of coverage by HDX-MS. White indicates regions of no peptide coverage by HDX-MS. Head and body domains, along with RNA binding loop and tail loop of NP are annotated based on Ye *et al*. (86).

### Ambient acidic exposure induces C-terminal flexibility in internal protein Matrix 1 (M1)

Lastly, we sought to determine whether the major capsid protein, M1, was structurally altered in pH-treated samples. M1 is an α-helical protein with two major domains; a globular N-terminal domain (NTD), and a comparably unstructured C-terminal domain (CTD), which are connected by a coiled linker sequence (54,55). M1 proteins form a ‘shell’ layer underneath the viral envelope, and exposure of M1 to low pH during virus entry is thought to cause the M1 layer to dissociate, ultimately allowing viral disassembly and genome release into the host cell cytoplasm (56–58). However, the M1 conformational changes that accompany this event are substantially less well characterized than those of HA.

HDX-MS analysis of M1 under neutral conditions (pH 7) revealed a highly dynamic CTD, indicated by multiple peptides showing >50% H/D exchange (**Fig. 6A**). Flexibility of the M1 C-terminus is in line with the current literature on M1, and has been reported to cause issues with resolution of this region using the standard and far more labor-intensive crystallographic methods (59). As with HA, the close membrane association of the N-terminal region of the M1 protein caused some coverage limitations, with substantially less peptides generated from the membrane binding domain for HDX-MS. Despite this, 68% of the overall M1 protein sequence was covered by at least one unique peptide (**Supp. Fig. S6**). When comparing pH 4-treated samples to this neutral state, H/D exchange rates of detected peptides in the M1 NTD were unaffected by pH treatment, with one exception (**Fig. 6B**). The loop encompassing residues 128-141 surprisingly showed protection from H/D exchange following pH 4 exposure (Peptide 1 in **Fig. 6C – D**). This was shown for two overlapping peptides covering the same region (**Fig. 6C**), and this observation suggests that this region adopts a more structured conformation upon pH 4 exposure. In agreement with this, prior work comparing X-ray crystal structures of M1 at neutral and acidic conditions indicated that structural differences were negligible within the NTD, apart from a loop centered around residue 135 (54). Again, this demonstrates the benefits of HDX-MS for structural analysis that is far less labor intensive than typical methods, whilst also enabling the virus to be retained in its whole native conformation.

**Figure 6.**
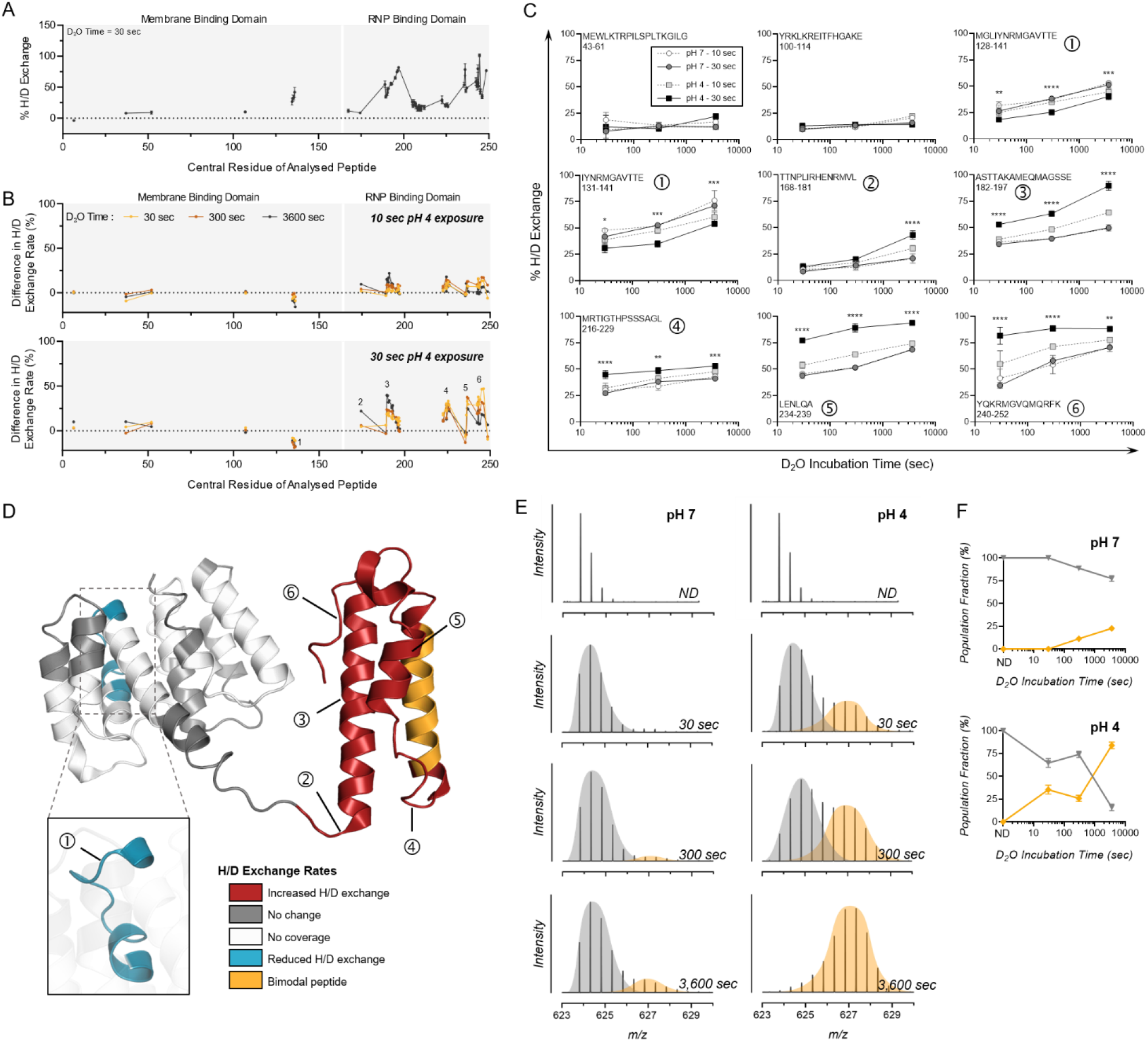
HDX-MS characterization of M1 protein from whole IAV after transient exposure to acidic conditions. **A)** H/D exchange percentages measured for M1 protein from whole IAV samples held at neutral conditions (pH 7) for 30 seconds. Each point shows the mean percentage of H/D exchange ± SD for a peptide represented by its central amino acid number (n=3, Uniprot: P05777). Continuous lines indicate regions of continuous peptide coverage by MS. Membrane binding domain and ribonucleoprotein (RNP) binding domain are indicated by grey shading and white lines. **B)** Data presented as residual plots showing the difference in H/D exchange percentages induced by exposure to pH 4 for 10 seconds or 30 seconds, when compared to corresponding pH 7 control samples. Peptides showing bimodal activity are excluded, unimodal peptides only are presented in panel B. Individual data points indicate the mean percentage of H/D exchange difference for triplicate samples. Peptides of interest showing increased H/D exchange are numbered 1-6. **C)** Uptake plots for selected Peptides 1-6 showing increased H/D exchange after pH 4 exposure. Data shows the total percentage of H/D exchange for pH 7 or 4 samples after D_2_O incubation for 30, 300, or 3600 seconds. The sequence and residue positions of each peptide are indicated in each panel. Data shows points indicate the mean percentage of H/D exchange ± SD for triplicate samples. Data analyzed by Two-way ANOVA (* p< 0.05, ** p < 0.01, *** p < 0.001, **** p < 0.0001, when comparing 30 second pH -exposed samples at each D_2_O time-point. Statistical results for comparison of 10 second pH-exposed samples appear in **Supplementary Table S2**). **D)** Protein regions showing increased (red) or decreased (blue) H/D exchange rates, or showing no change in H/D exchange rates (grey) after 30 seconds at pH 4 (compared to pH 7 control) mapped onto the crystal structure of A/PR8 M1 (PDB: 7JM3). White indicates regions of no peptide coverage, peptides with bimodal activity are colored orange. Location of Peptides 1-6 from Panel C are also indicated by numbering. **E)** Representative spectra of triplicate samples from pH 7 and pH 4 groups showing a mass shift for bimodal peptide 204-218 with increasing D_2_O incubation times. Bimodal peaks are separated by grey and orange coloring. **F)** Proportions of the left (grey) and right (orange) peaks are graphed with time. Data indicates the mean ± SD for triplicate samples analyzed by HDX-MS.

In sharp contrast to the ordered NTD, the CTD encompassing alpha-helices H10 to H12 exhibited significantly increased H/D exchange rates post-pH 4 exposure (**Fig. 6B – D**). These results also correlate well with past work using small-angle X-ray scattering and atomic force microscopy, which suggested that a low-pH induced conformational change in M1 was most pronounced in the CTD and in the linker region between N and CTDs (59). Several peptides encompassing residues 202-212 in helix H11 also showed a bimodal distribution at neutral conditions, which was significantly more pronounced following pH 4 exposure (**Fig. 6E – F**). This H/D exchange regime is reminiscent of long-range conformational changes and suggest that exposure to acidic treatment promotes transition of an alpha-helix to a loop structure. The absence of peptides showing EX1 kinetics in helix H10 and H12, in contrast with neighboring helix H11, may be explained by the more structured conformation of the latter helix. Indeed, helices H10 and H12 have very high deuteration levels after only 30 seconds deuteration, whereas helix H11 is more protected from exchange even after 1-hour incubation in deuterated buffer (**Supp. Table S5**). The high magnitude of differences in H/D exchange rates observed for the CTD suggest a large unfolding of the alpha-helical structures upon acidic pH treatment, which may disrupt intra-molecular contacts known to exist between the CTD and NTD of neighboring M1 molecules (55) and may lead to partial disassembly of the M1 shell.

## DISCUSSION

Environmental transmission relies on viruses staying infectious long enough to come into contact with the next host. Non-enveloped viruses are typically considered more stable than enveloped viruses (60,61), particularly in aqueous environments (62). Despite this sensitivity, many enveloped viruses do undergo effective environmental transmission, with expelled liquid droplets and aerosols being major transmission vectors. Low droplet pH has been shown to inactivate enveloped phi6 bacteriophage during fomite transmission (63), and our prior work identified low pH as a major factor driving inactivation of IAV in aerosol particles (29). This was supported by infectivity data here (**Fig. 1B**), which showed >5-log_10_ decrease in infectious IAV titers within 30 seconds of low pH exposure. Small aerosol particles rapidly acidify in indoor air, though acidification is predicted to be slower for larger aerosol particles (**Fig. 1A**). Thus it may be that larger aerosol particles are the predominant carriers of infectious IAV. However, studies indicate that IAV is likely carried within the small aerosol size fraction, with both respiratory viral genomes and infectious IAV found to be enriched in fine aerosol particles ≤ 5 μm compared with coarser aerosols in humans (23,64). Thus IAV is likely carried within small aerosols, and would quickly be exposed to acidic conditions upon aerosol exhalation.

In the present study, we sought to improve understanding of the effects that this acidic microenvironment may exert on the structure of IAV during environmental transmission. We placed particular focus on the effects of low pH on structural lipid, genome, and protein components of IAV, to investigate the processes mediating rapid virus inactivation. Understanding the mechanisms leading to aerosol-associated pathogen inactivation will be useful for public health, as these pathways could be targeted by novel treatments to enhance pathogen inactivation in the air. All experiments were performed with whole virus rather than purified molecules to mimic the natural virus setting where many stabilizing protein-protein and protein-lipid interactions are present. Collectively, our whole-virus data showed that viral proteins rather than lipids or genome were undergoing measurable changes over the time-course relevant to the observed pH-mediated viral decay. Multiplexed whole-virus HDX-MS data pointed to both HA (**Fig. 1 – 2**) and M1 (**Fig. 6**) proteins as drivers of virus inactivation. The remaining structural proteins of IAV (M2, NA) and the replication associated proteins (PA, PB1, PB2) were at too low of a copy number per virion to analyze using our whole-virus HDX-MS method. Viral ribonucleoprotein has previously been found to aggregate at low pH (∼5), though this aggregation was mostly reversed by pH neutralization (65) suggesting no permanent changes occur. The amplified region of the IAV genome was also unaffected in our study by low pH exposure (**Supp. Fig. S3**), and NP, which coats the IAV genome segments, showed no pH-induced conformational changes by HDX-MS analysis (**Fig. 5B – C**). Thus, structural damage to replication-associated components of IAV are unlikely to be contributing to the loss of viral infectivity observed in acidic conditions.

Functionality data also confirmed the premature attainment of the post-fusion form of HA as a primary cause of rapid IAV inactivation under acidic aerosol conditions (**Fig. 1C – E**). The pH-mediated change of HA is well-documented in the literature, however conditions described to trigger this involve longer and hotter incubations at pH ∼5.5. Despite the very rapid time scale, the ambient temperature, and the substantially lower pH used here (1.5 pH units below fusion), HA was still shown to transition to the post-fusion state, or at the very least was triggered to enter a stable intermediate state that prevents viral attachment and entry. Other studies have reported intermediate HA states between pre- and post-fusion (31,43,66–69), some of which seem to also be reversible (70–72). The retained flexibility and bimodal H/D exchange behavior of the β-loop in particular (**Fig. 3A – C**) indicated that for our pH-exposed samples, HA was in an intermediate state, similar to that shown by Benhaim *et al*. (43) for H3N2 HA in early points post-pH exposure. Importantly, re-neutralization of our samples prior to any HDX-MS analysis indicated an irreversible transition had occurred, leading to definitive loss of viral infectivity after acidification. Additional experiments testing longer exposure times or higher temperatures (37°C as opposed to ambient) would help in classifying the degree of flexibility in the β-loop post-pH exposure in our system.

Interestingly, changes detected in HA at 10 seconds were not amplified by 30 seconds (**Fig. 2C**), despite viral titers continuing to drop (**Fig. 1B**). Thus, whilst HA appeared to mediate the majority of inactivation, changes to M1 and the associated capsid disassembly could drive additional infectious titer loss. In fact, inactivation kinetics in **Figure 1B** could be classified as two-step kinetics, with a sharp initial drop in infectivity from 0 to 10 seconds post-pH exposure, followed by a slower rate of decay up to 30 seconds. This second portion of inactivation could be attributed to M1 conformational changes and associated capsid disassembly. In cryo-EM studies, incubation of H3N2 IAV at low pH was reported to induce loss of spherical virion morphology and a disruption and/or coiling of the matrix layer away from the viral envelope in some virions, whilst other virions in the same suspension showed unaffected morphology and a resolvable matrix later (56,70). This suggests M1 disassembly affects some but not all virions in a population at the same rate. This was similarly observed for H1N1 A/WSN/33 here, with some but not all virions showing loss of spherical morphology by TEM after low pH treatment (**Fig. 4A**). The virion integrity assay (**Fig. 4B**) also indicated that lysed viral particles could account for ∼2-log_10_ in viral titer reduction by 30 seconds post-pH exposure. This would supplement the initial sharp 3-log_10_ reduction seen over the first 10 seconds, which we attribute to HA changes. Whilst the first ∼30 seconds of acid exposure was of the most interest here, it could also be beneficial to classify the state of these IAV proteins after longer exposure times (e.g. 5 minutes, 30 minutes) in a subsequent study, to assess ongoing changes.

We acknowledge the limitations of this study for determining the effect of low aerosol pH on IAV in combination with other aerosol factors, including the presence of protective proteins and other organic components. We recognize that the salt-based pH buffers used here do not reflect the complexity and variability of lung and nasal fluids from human patients, in which IAV would be suspended in a physiological aerosol setting, and these may offer partial protection and reduce acid-mediated conformational changes to HA and/or M1. For example, IAV has been demonstrated to readily bind sialic acids present on respiratory mucins (73–75). These mucins are part of the protective mucus layer secreted by epithelial cells of the respiratory tract, and presentation of sialylated glycans by respiratory mucins is intended to act as a decoy receptor for the virus (75,76). This IAV-mucin interaction has also been shown to be weaker than the true IAV-host cell interaction (77) and is often reversible due to the action of viral neuraminidase (73,78). It is therefore feasible that a transient interaction such as this could be stabilizing for the virus during aerosol transport between hosts. In fact, Hirose *et al*. demonstrated IAV suspended in simulated gastric acid (pH 2) rapidly lost infectivity, but could be protected from acid-mediated inactivation for at least 4 hours if co-incubated with artificial mucus (79). It will be the focus of a follow-up study to determine if mucins can mediate stabilization of IAV proteins through aerosol-associated stress. However, our prior work has shown that protective effects of multiple respiratory matrices are no longer functional in bulk solution at a lower pH of 4 (29), thus their inclusion would not have altered the outcomes of this particular study. Additionally, the aim of the current study was to intentionally decouple the effect of low pH from other factors (whether inactivating or protective) within an evaporating aerosol, for mechanistic characterization. The pH sensitivity of the A/WSN/33 strain used here is also similar to that of other strains, both lab-adapted and clinical isolates (80), and WSN was intentionally used as a surrogate as clinical strains are typically harder to work with and don’t grow to the high titers required for HDX-MS analysis. Regardless, the panel of strains tested could be expanded in future work to gain additional sensitivity information.

Beyond the scope of this study, it will also be of interest to determine whether other respiratory viruses (e.g. MERS CoV, rhinovirus, respiratory syncytial virus, etc.) will be similarly affected by acidic aerosol pH, and whether a broad acting air treatment could be employed as a non-pharmaceutical intervention to reduce infectious viral loads in the air. Alternatively, a spray based therapeutic could be utilized to accelerate the acidification of exhaled aerosols in high-risk settings, enhancing the speed of viral inactivation. A prior study investigated the efficacy of a low pH (∼3.5) nasal spray for mitigating the effects of IAV infection in a ferret model. The nasal sprays tested were based on simple acid mixtures (including L-pyroglutamic acid, succinic acid, citric acid and ascorbic acid), and their topical application to infected ferrets reduced severity of viral-induced symptoms and reduced viral shedding (81). As multiple epitopes on HA were found to be affected by pH here, the non-specificity of low pH to drive IAV inactivation means that this approach may be less prone to resistance development than current antiviral drugs. IAV is also specifically reliant upon acidic pH exposure for progression of its natural infection cycle *in vivo*, thus development of resistance to acidic treatments is highly unlikely.

Overall, information gained here improves our understanding of the innate inactivation mechanisms for IAV under indoor aerosol conditions. Whole-virus HDX-MS has only been used previously to characterize the HA protein of H3N2 IAV strain X-31. We determined that IAV was not inactivated due to global protein denaturation, but instead have confirmed that localized structural changes occur to HA under aerosol-associated pH 4 conditions, similar to those changes induced by the acidic host cell endosome. We have also identified an additional mechanism of M1-driven inactivation. Expanding the existing body of IAV-specific HDX-MS data to encompass additional influenza virus strains and subtypes will be useful for the field in general to enable study of mechanistic similarities/differences between viral proteins, e.g. in terms of pH susceptibilities of different HA and M1 molecules. This could be used to predict environmental stability and transmission risks of emerging strains, assess the suitability of air treatment strategies, or to perform comparison of epitope flexibilities to predict the efficacy of neutralizing antibodies or antivirals.

## MATERIALS & METHODS

### Virus Stocks

Influenza A virus (IAV) strain A/WSN/33 [H1N1 subtype] was propagated in Madin-Darby Canine Kidney (MDCK) cells (ThermoFisher). Cells were maintained in Dulbecco’s modified Eagle’s medium (DMEM, Gibco) supplemented with 10% Fetal Bovine Serum (FBS; Gibco), and 1% Penicillin-Streptomycin 10,000 U/mL (P/S; Gibco). Confluent MDCK monolayers were washed and inoculated with IAV at a MOI of 0.001 for 72h in OptiMEM (Gibco) supplemented with 1% P/S and 1 μg/mL TPCK trypsin (Sigma, T1426). Infected culture supernatants were clarified by centrifugation at 2,500 ×*g* for 10 min, and IAV was then pelleted through a 30% sucrose cushion at 112,400 ×*g* in a SW31Ti rotor (Beckman) or an AH-629 rotor (ThermoFisher Scientific) for 90 min at 4°C. Pellets were recovered in phosphate buffered saline (PBS, ThermoFisher, 18912014) overnight at 4°C. Concentrated IAV stocks were quantified at 1.2 × 10^10^ to 7.5 × 10^11^ PFU/mL by plaque assay (described below), and at 5 mg/mL total viral protein by Qubit Protein Assay quantification kit (Life Technologies, Q33211).

Recombinant Vesicular Stomatitis Virus (VSV) stocks harboring a Firefly Luciferase gene instead of VSV G (Kerafast, EH1020-PM) were pseudotyped with either the VSV G protein (VSV-G) or the A/WSN/1933 HA and NA (VSV-H1N1). 8 × 10^5^ HEK293T cells/well were seeded into 6-well plates and transfected the next day with ViaFect (Promega, E4981), using either 4 μg of plasmid expressing the G protein or 2 μg of plasmid expressing HA together with 2 μg of plasmid expressing NA. After incubation at 37°C for 24 h, cells were infected with a previously made VSV-G Luc stock at a MOI of 4 in DMEM. After 1.5 h at 37°C cells were washed with PBS and incubated with DMEM containing 1 μg/ml anti-VSV-G antibody (Kerafast, EB0010) for 24 h at 37°C. Note that this was added to the VSV-H1N1 wells only, VSV-G stocks were generated without the antibody. Supernatant was collected and clarified by centrifugation at 300 ×*g* for 5 min. VSV-H1N1 stocks were additionally treated with 5 μg/mL TPCK-treated trypsin (Sigma-Aldrich, T1426) for 20 min at 37°C, followed by trypsin inactivation with 10 μg/ml Soybean Trypsin inhibitor (Sigma-Aldrich, T6522) for 20 min at 37°C. VSV-G and VSV-H1N1 stocks were aliquoted and frozen at -80°C.

### Plaque Assay

Plaque assay was conducted using monolayers of MDCK cells in 12-well plates. IAV samples were serially diluted in ‘PBS for infections’ (PBSi; PBS supplemented with 1% P/S, 0.02 mM Mg^2+^, 0.01 mM Ca^2+^, 0.3% bovine serum albumin [BSA, Sigma-Aldrich A1595], final pH ∼7.3), before being added to washed cellular monolayers. Viruses were incubated on monolayers for 1 h at 37°C with 5% CO_2_ with manual agitation every 10 min. Non-attached viruses were removed from cells by washing with PBS, and MEM supplemented with 0.5 μg/mL TPCK-trypsin and agarose was added to cells. Infected and control plates were incubated for 72 h at 37°C with 5% CO_2_, and plaques were visualized after fixing cells in PBS + 10% formaldehyde (Sigma, 47608-1L-F), then staining with 0.2% crystal violet solution (Sigma, HT901-8FOZ) in water + 10% methanol (Fisher Chemical, M-4000-15).

### Virus Acidification & Neutralization

Citric acid powder (Acros Organics, 220345000) and Na_2_HPO_4_ powder (Fluka Chemika, 71640) were dissolved in miliQ H_2_O to reach concentrations of 0.1 M and 0.2 M respectively. 0.1 M citric acid and 0.2 M Na_2_HPO_4_ solutions were then mixed together as specified in the Sigma Aldrich Reference System (accessed here: https://www.sigmaaldrich.com/CH/en/technical-documents/protocol/protein-biology/protein-concentration-and-buffer-exchange/buffer-reference-center#citric) to generate 1× pH 4 or 7 salt-based buffers. Alternatively, concentrated 1M citric acid and 2 M Na_2_HPO_4_ solutions were made, and mixed in the same ratios to generate 10× pH 4 or 7 buffers. For IAV infectivity analysis, 1× pH buffers were used to pH adjust virus samples. Purified IAV was spiked ∼1/100 into 1× pH 4 or pH 7 buffers (final concentration 1 × 10^7^ PFU/mL or 1 × 10^8^ PFU/mL), vortexed briefly to mix, and left to incubate at ambient temperature. At specified time-points, samples were taken and diluted 1/100 in PBSi to neutralize pH. Alternatively, samples were neutralized by a 1/10 dilution in PBSi containing 3% 10× pH 7 buffer to minimize virus dilution. pH testing ensured both dilution methods neutralized the pH to 6.7 – 7.0 in all cases. Neutralized samples were frozen prior to titration by plaque assay or fluorescence assay for quantification of remaining infectious titers.

For HDX-MS, lipid-MS, VSV analyses, TEM imaging, and RNase Capsid Integrity Assay, 10× pH buffers were used to adjust purified virus samples to pH 4 or 7, by adding 10× pH buffers to the virus sample in a 1:9 ratio. This strategy allowed pH to be finely tuned whilst keeping virus sample concentrations as high as possible. For example, to adjust to pH 4, 1 μL of acidification buffer (10× pH 4.0 buffer, comprised of 600 mM citric acid, 800 mM Na_2_HPO_4_) was added to 9 μL of virus sample, and vortexed for 2 seconds to mix. This reduces sample pH from ∼7.3 to 4.0. To neutralize IAV samples, 2.2 μL of 2M Na_2_HPO_4_ was added to the 10 μL mixture of acidified virus, and vortexed for 2 seconds to mix. Note that mixing non-adjusted virus with 2M Na_2_HPO_4_ in the same ratio did not impact IAV infectious titers (**Supp. Fig. S7**). To neutralize VSV samples, acidified VSV was diluted 1/10 in PBSi containing 3% 10× pH 7 buffer. Both neutralization methods restored pH to a neutral pH of 6.7 to 7.0. For neutral control samples, 1 μL of 10× pH 7.2 buffer (160 mM citric acid, 1.68 M Na_2_HPO_4_) was mixed with 9 μL of virus sample and vortexed for 2 seconds to mix, as a mock acidification. This reduced sample pH from ∼7.3 to 7.0. To mock neutralize, 2.2 μL of 10× pH 7.0 buffer (130.5 mM citric acid, 1.74 M Na_2_HPO_4_) was added to this 10 μL and vortexed for 2 seconds to mix. This retained pH at 7.0, and this strategy ensured salt concentrations and virus concentrations were comparable between neutralized pH 4 and pH 7 samples prior to downstream analysis. All pH adjustments and incubations were performed at ambient room temperature (22 – 25°C).

### VSV Infectivity Analysis

To measure inactivation of pseudotyped VSV, samples were acidified and neutralized as described above and subsequently tested for entry capacity by measuring the luciferase activity of infected MDCK cells. To do so, 3 × 10^4^ cells/well were seeded into 96-well plates and incubated overnight. The next day, cells were washed and infected with 50 μL of neutralized VSV sample for 1.5 h at 37°C. Cells were washed with PBS and incubated in DMEM with 10% FBS and 1% P/S. Luciferase activity was quantified 7 hours post infection using the ONE-Glo Luciferase Assay System (Promega, E6110) in combination with the PerkinElmer EnVision plate reader.

### Fluorescence Imaging of Infected Cells & Viral Particles

IAV samples were acidified and neutralized as described above, resulting in final concentrations of 1 × 10^7^ PFU/mL. Samples were then used for infection and subsequent staining of intracellular NP or direct staining of viral particles. To evaluate virus entry after acidification, A549 cells were infected for 1 h at 37°C with neutralized IAV diluted in PBSi at a multiplicity of infection (MOI) of 2 (calculated from titers prior to acidification). Afterwards, cells were washed with PBS and incubated for 2 or 5 h in DMEM supplemented with P/S, 0.2% BSA, 20 mM HEPES (Sigma-Aldrich, H7523) and 0.1% FBS. Infected cells were fixed for 15 min with 3.7% formaldehyde (ThermoFisher Scientific, 26908) in PBS and subsequently blocked and permeabilized for 1 h at RT in PBS supplemented with 50 mM ammonium chloride (Sigma-Aldrich, 254134), 0.1% saponin (Sigma-Aldrich, 47036), and 2% BSA (Sigma-Aldrich, A7906). Primary murine monoclonal anti-NP antibody (hybridoma supernatant, ATCC, HB-65) was applied for 1 h at RT (or overnight at 4°C when required). Cells were washed in PBS, and bound primary antibody was detected using anti-mouse IgG Alexa488 (ThermoFisher Scientific, A-11029). Infected cells were also stained with DAPI (Sigma-Aldrich, 10236276001) and incubated at RT for 1 h.

To analyze HA conformation of viral particles, 150 μL of neutralized IAV was centrifuged onto cover slips for 30 min at 1000 ×*g*. Virus samples were then fixed, blocked, and permeabilized as described above, and IAV was stained using a murine monoclonal post-fusion HA antibody (hybridoma supernatant, Wistar - Coriell Institute, Y8-10C2, validated in (82) to bind more strongly to post-fusion HA), and a rabbit polyclonal anti-NP antibody (kind gift of J. Pavlovic, Institute of Medical Virology, Zurich, Switzerland). Staining for internal NP ensured the same amount of viral material was imaged in all cases. Stained samples were washed in PBS, and bound antibodies were detected using anti-mouse IgG Alexa546 (ThermoFisher Scientific, A-11031) and anti-rabbit IgG Alexa488 (ThermoFisher Scientific, A-11008). Following washing in PBS, cover slips were mounted with ProLong Gold Antifade Mountant (Thermo Fisher Scientific, P36930). Stained slides and cells were imaged with the SP8 confocal laser scanning microscope (Leica) or the DMi8 microscope (Leica) in combination with the THUNDER Instant Computational Clearing algorithm (Leica).

### Transmission Electron Microscopy

Purified IAV samples were pH-adjusted and neutralized using 10× pH buffers as described above. All samples were pH exposed for 30 seconds at ambient room temperature prior to neutralization. Neutralized samples were then diluted to 10^10^ PFU/mL equivalent in 1× pH 7.0 buffer and fixed with 0.06% paraformaldehyde for 4 days at 4°C prior to use in TEM. 5 μL of each fixed sample was deposited onto carbon-coated copper grids (Electron Microscopy Sciences, CF400-Cu). Grids were UV-glow discharged immediately prior to sample addition, and sample was left to adsorb for 2 min at room temperature. Excess solution was blotted using Whatman filter paper, and grids were washed in a droplet of miliQ water. Samples were then negatively stained for 1 min with 5 μL of 2% uranyl acetate (Electron Microscopy Sciences, 541-09-3. Excess stain was blotted, and grids were allowed to air dry for 5 min. Grids were imaged using the Tecnai Spirit TEM (spot size: 2, emission: 4.41 uA, accelerating voltage: 80 kV).

### RNase Assay for Capsid Integrity

IAV samples were pH-adjusted and neutralized as described above, using 1× pH buffers. Samples were pH exposed for 30 seconds at ambient room temperature prior to neutralization. As a control, virus was mixed with PBS only and held at ambient temperature. Neutralized and control whole-virus samples were then diluted 1/100 into RNase digestion mixture (TE buffer with 10 mM Tris and 1 mM EDTA [Invitrogen, AM9858], supplemented with 50 mM NaCl, pH adjusted to 7.0 and with HCl. RNase A/T1 [Thermo-Scientific, EN0551] then added for 40 μg/mL RNase A and 100 U/mL T1 final concentrations). Samples vortexed to mix, and statically incubated at 37°C for 30 min. As controls, identical neutralized samples were diluted 1/100 into control digestion mixture (PBS added to TE buffer in the absence of RNase A/T1), vortexed, and incubated alongside RNase-treated samples. After 30 min, SUPERase-In RNase Inhibitor (Invitrogen, AM2694, acts against RNase A, B, C, 1, T1) was added to all samples, and incubated at room temperature for 20 min Samples were then frozen at -20°C overnight prior to RNA extraction using the QIAamp Viral RNA Mini extraction kit (Qiagen, 52906) according to manufacturer’s instructions. Viral RNA was stored at -80°C until analysis.

### RT-qPCR

Amplification and detection were performed using the One Step PrimeScript™ RT-PCR Kit (RR064A, Takara Bio) with the following primers targeting the IAV M segment: forward primer 5’-ATGAGYCTTYTAACCGAGGTCGAAACG-3’ and reverse primer 5’-TGGACAAANCGTCTACGCTGCAG-3’. The RT-qPCR mixture for IAV detection was 7.5 μL of 2× One-Step SYBR RT-PCR Buffer, 0.3 μL of Takara Ex Taq HS (5 U/μL stock), 0.3 μL PrimeScript RT enzyme mix, 0.3 μL forward and reverse primers (10 uM stocks), 3.3 μL RNase-free water, and 3 μL extracted RNA sample. All sample manipulations were performed under biosafety cabinets, and the following conditions were used for the one-step cycling program: 5 min at 95°C for denaturation, followed by 40 cycles of 95°C for 5 sec, 60°C for 20 sec, and 72°C for 10 sec for annealing and extension. For quantification, we created a standard curve using a titration of a Gblock gene fragment, as described in (83). All RT-qPCR reactions were performed in duplicate, and each PCR run contained non-template controls which were always negative. Sample dilutions ensured no PCR inhibition in our experimental protocol. Reactions were measured using a Mic Real-Time PCR System from Bio Molecular Systems.

### ResAM Modelling

The multi-layer Respiratory Aerosol Model (ResAM) simulates the composition and pH changes inside an expiratory particle following exhalation and subsequent equilibration with indoor air (for a detailed description of the viral study with ResAM, see (29)). The model performs calculations for particles of selectable size (from 20 nm to 1 mm) with a liquid phase composed of organic and inorganic species representative of human respiratory fluids, specifically synthetic lung fluid. Conditions of indoor room air were set at 20°C and 50% relative humidity, with concentrations of 36.3 ppb NH_3_, 0.27 ppb HNO_3_, and 600 ppm CO_2_, as described in the above study.

### Hydrogen-Deuterium exchange coupled to Mass-Spectrometry (HDX-MS)

Purified IAV samples were pH-adjusted and neutralized as described above with 10× pH buffers. Samples were pH exposed for 10 or 30 seconds at ambient room temperature prior to neutralization. Careful tests were run to make sure acid-treated and pH 7-treated samples had the exact same pH following neutralization to avoid differences in hydrogen/deuterium exchange (HDX) rate due to shifts in pH during deuteration. To initiate HDX, neutralized whole-virus samples were incubated in D_2_O buffer (99% deuterium oxide, Sigma 151882, supplemented with 120 mM NaCl). 40 μL D_2_O buffer was added to every 10 μL of neutralized virus (final D_2_O concentration 78%) on ice for 30 seconds, 300 seconds, or 3,600 seconds in triplicate. After each D_2_O incubation, samples were quenched by dilution into ice-cold quench buffer (miliQ containing 150 mM tris(2-carboxyethyl) phosphine-HCl [TCEP, Thermo-Scientific, PG82080], 2 M urea [Sigma, U5128], and 0.1% formic acid [FA, Acros Organics, 270480280] final concentrations, pH 2.5). Samples were spun at 21,000 ×*g* for 2 min in a pre-cooled 4°C centrifuge, and supernatant was immediately flash-frozen in liquid nitrogen then stored at -80°C until analysis by HDX-MS. Non-deuterated (ND) controls were prepared identically using pH 7 and pH 4 neutralized virus, using H_2_O supplemented with 120 mM NaCl instead of D_2_O.

### Deuterium incorporation quantification

To quantify deuterium uptake into the protein, protein samples were thawed and injected in UPLC system immersed in ice. The protein was digested via two immobilized pepsin columns (Thermo, 23131), and peptides were collected onto a VanGuard precolumn trap (Waters). The trap was subsequently eluted and peptides separated with a C18, 300Å, 1.7 μm particle size Fortis Bio column 100 × 2.1 mm over a gradient of 8 – 30 % buffer B over 20 min at 150 μL/min (Buffer A: 0.1% formic acid; buffer B: 100% acetonitrile). Mass spectra were acquired on an Orbitrap Velos Pro (Thermo), for ions from 400 to 2200 m/z using an electrospray ionization source operated at 300 °C, 5 kV of ion spray voltage. Peptides were identified by data-dependent acquisition after MS/MS and data were analyzed by Mascot. Deuterium incorporation levels were quantified using HD examiner software (Sierra Analytics), and quality of every peptide that was checked manually. Results are presented as percentage of theoretical maximal deuteration levels. Extended results are found in Supplementary Tables S3, S4 and S5. Changes in deuteration level between two states were considered significant if >12% and >0.6Da and p<0.05 (unpaired t-test).

### Whole Virus Lipid Mass Spectrometry

Purified IAV samples were diluted to 8 × 10^10^ PFU/mL in PBS, then pH-adjusted and neutralized after 30 seconds of pH exposure at ambient temperature, as described above. Neutralized whole-virus samples were then diluted 1/100 in chloroform:methanol (1:1 mixture) using glass pipettes, in 2 mL Eppendorf Safe-Lock Polypropylene tubes. Samples were briefly vortexed to mix, and kept at 4°C until analysis. Lipid extracts were separated over an 8-minute gradient at a flow rate of 200 μL/min on a HILIC Kinetex Column (2.6lm, 2.1 × 50 mm^2^) on a Shimadzu Prominence UFPLC xr system (Tokyo, Japan). Mobile phase A was acetonitrile:methanol 10:1 (v/v) containing 10 mM ammonium formate and 0.5% formic acid, while mobile phase B was deionized water containing 10 mM ammonium formate and 0.5% formic acid. The elution gradient began at 5% B at 200 μL/min and increased linearly to 50% B over 7 min, held at 50% B for 1.5 min, and finally, the column was re-equilibrated for 2.5 min. MS data were acquired in full-scan mode at high resolution on a hybrid Orbitrap Elite (Thermo Fisher Scientific, Bremen, Germany). The system was operated at 240,000 resolution (*m/z* 400) with an AGC set at 1.0E6 and one microscan set at 10-ms maximum injection time. The heated electrospray source HESI II was operated in positive mode at 90 °C and a source voltage at 4.0KV. Sheath gas and auxiliary gas were set at 20 and 5 arbitrary units, respectively, while the transfer capillary temperature was set to 275 °C.

Mass spectrometry data were acquired with LTQ Tuneplus2.7SP2 and treated with Xcalibur 4.0QF2 (Thermo Fisher Scientific). Lipid identification was carried out with Lipid Data Analyzer II (LDA v. 2.6.3, IGB-TUG Graz University) (84). The LDA algorithm identifies peaks by their respective retention time, *m/z* and intensity. Care was taken to calibrate the instrument regularly to ensure a mass accuracy consistently lower than 3 ppm thereby leaving only few theoretical possibilities for elemental assignment. Data visualization was improved with LCMSexplorer in a homemade web tool hosted at EPFL.

### Statistical Analysis

Quantitative results expressed as mean ± SD. Student’s t-test was used for comparison of data from 2 groups, One-way ANOVA used for comparison of data from 3 or more groups involving a single independent variable, Two-way ANOVA used when data was grouped according to two independent variables. Tukey’s multiple comparisons test used for post-hoc analysis. All analyses performed using GraphPad Prism, version 9.4.0 (GraphPad Software, La Jolla, USA). P-values <0.05 (95% confidence) considered statistically significant.

## Acknowledgements

This work was funded by the Swiss National Science Foundation (Grant #189939). We thank Rémy Visentin and Alexandre Hainard from the protein and proteomics platforms at the University of Geneva for assistance with HDX-MS sample acquisition. We thank Davide Demurtas of the EPFL Interdisciplinary Centre for Electron Microscopy (CIME) for assistance with TEM imaging.

## Data availability statement

All data relating to this study will be shared in accordance with FAIR (findable, accessible, interoperable, and reusable) data principles, and data will be deposited in the community-approved and cross-disciplinary public repository Zenodo, upon acceptance of publication. The mass spectrometry proteomics data have been deposited to the ProteomeXchange Consortium via the PRIDE partner repository (85) with the dataset identifier PXD037176.

