## Supplementary Material for "Inactivation mechanisms of Influenza A virus under pH conditions encountered in aerosol particles as revealed by whole-virus HDX-MS"

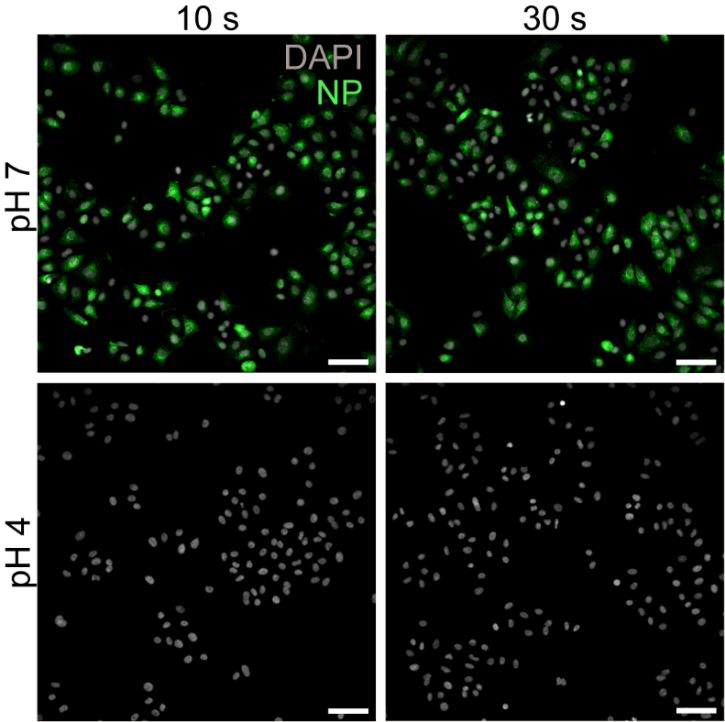

**Supplementary Figure S1 – Immunofluorescence images of A549 cells infected with IAV samples treated at pH 4 or 7 for 10 or 30 seconds.** All samples were neutralized prior to cell infections. A549 cells were infected at a multiplicity of infection (MOI) of 2, and IAV protein production was visualized by immunofluorescent staining of the nucleoprotein (NP) at 6 hours post-infection. Scale bar corresponds to 250 μm. Images are representative of duplicate experiments.

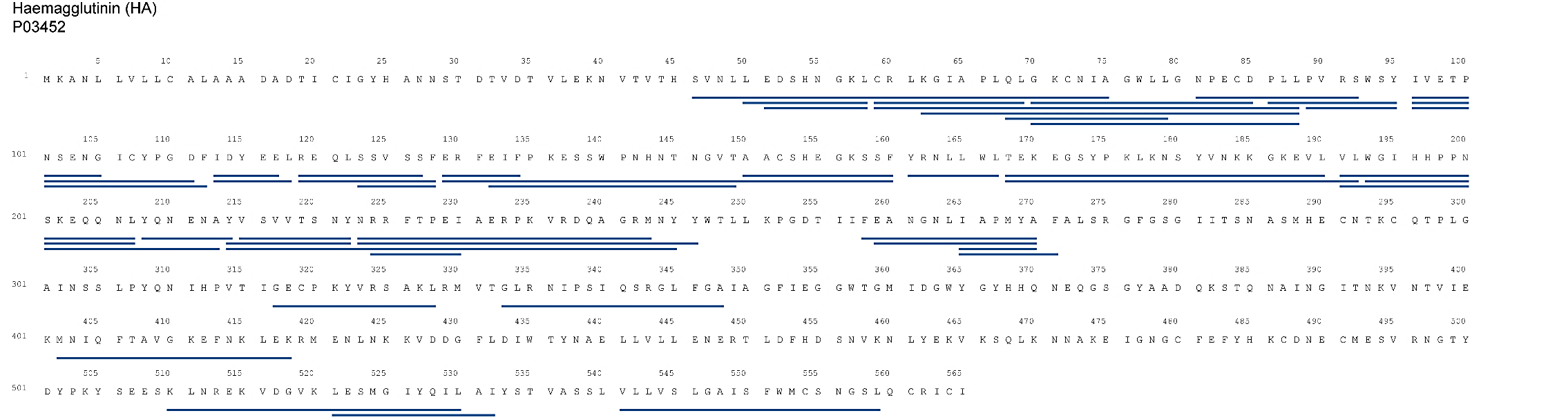

**Supplementary Figure S2** – **HDX-MS peptide coverage map displaying unique peptides for influenza A virus HA.** Whole virus samples were acidified to pH 4, then neutralized back to pH 7 after 30 seconds. Samples were then incubated in D_2_O, quenched and processed for analysis by HDX-MS. Peptides aligned to published HA protein sequence for strain A/PR8 (UniProt: P03452). Only unique peptides where HDX-MS data was attained for acid-treated samples are shown here. Under these conditions, 74% of the HA1 protein subunit (residues 18 – 343) and 30­% of the HA2 protein subunit (residues 344 – 565) were covered by at least 1 unique peptide. Coverage of HA2 was limited compared with HA1, which is attributed to protease resistance and steric occlusion from high HA2 density on intact IAV virions. Coverage maps made using HDExaminer (Sierra Analytics).

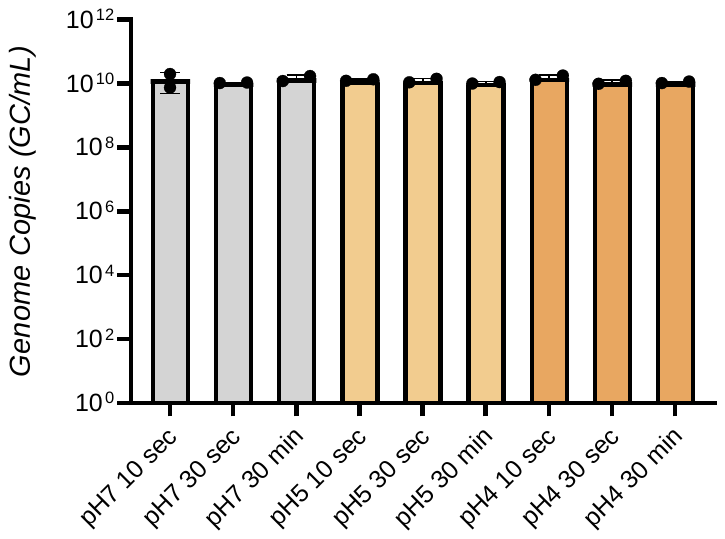

**Supplementary Figure S3 – Pre-treatment with acidic pH does not alter amplifiable copies of viral RNA, even after 30 minutes.** RT-qPCR results showing the IAV genome region tested (M2 genome segment) is able to be replicated to comparable levels regardless of pH 4, 5 or 7 pre-treatments for 10 seconds, 30 seconds, or 30 minutes at room temperature. Samples were all neutralized back to pH 7 prior to RNA extraction and quantification by RT-qPCR. Data presented as mean genome copies (GC/mL) ± SD for duplicate samples.

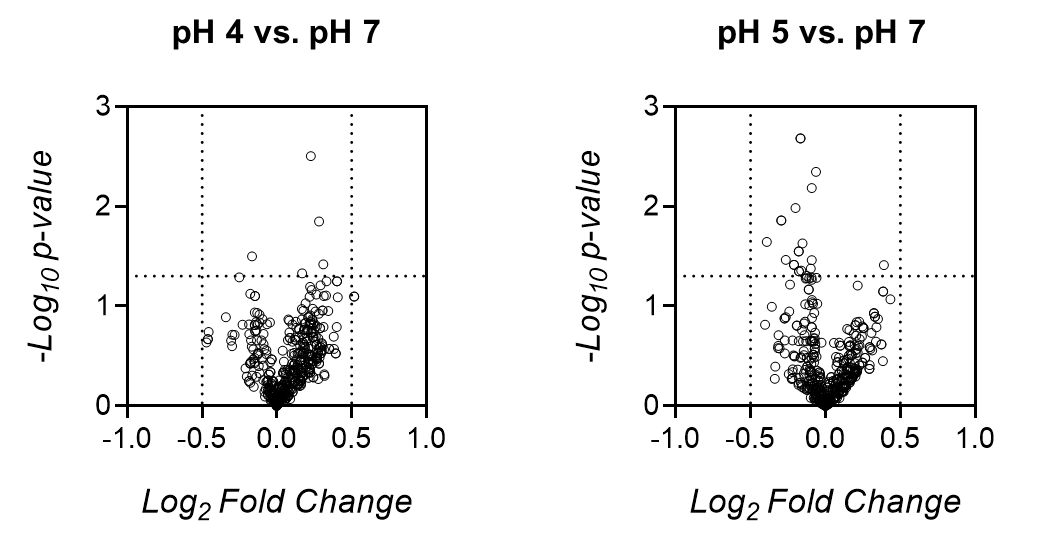

**Supplementary Figure S4 – Pre-treatment with acidic pH does not significantly alter lipid classes of the IAV envelope.** Volcano plots obtained by comparing the relative levels of lipid species in samples exposed to pH 4 (left panel) or pH 5 (right panel) to samples exposed to pH 7. N = 3 samples per treatment group, each data point indicates an individual lipid species, vertical dotted lines are drawn to highlight size effects > 0.5 or < -0.5 Log_2_-Fold change, horizontal dotted lines are drawn to highlight significant changes (-Log_10_ p-values > 1.301; i.e., p-value < 0.05)*.*

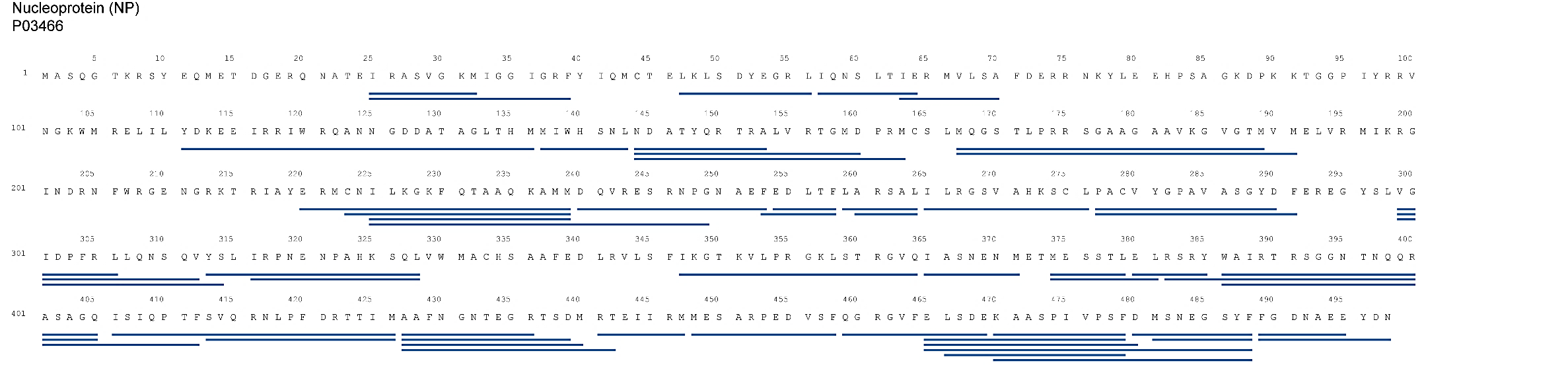

**Supplementary Figure S5 – HDX-MS peptide coverage map displaying unique peptides for influenza A virus NP.** Whole virus samples were acidified to pH 4, then neutralized back to pH 7 after 30 seconds. Samples were then incubated in D_2_O, quenched and processed for analysis by HDX-MS. Peptides aligned to published NP protein sequence for strain A/WSN/33 (UniProt: P03466). Only unique peptides where HDX-MS data was attained for acid-treated samples are shown here. In these conditions, 75% of the NP protein was covered by at least 1 unique peptide. Coverage was evenly distributed across the NP protein, with slightly more unique peptides appearing at the C-terminal end. Coverage maps made using HDExaminer (Sierra Analytics).

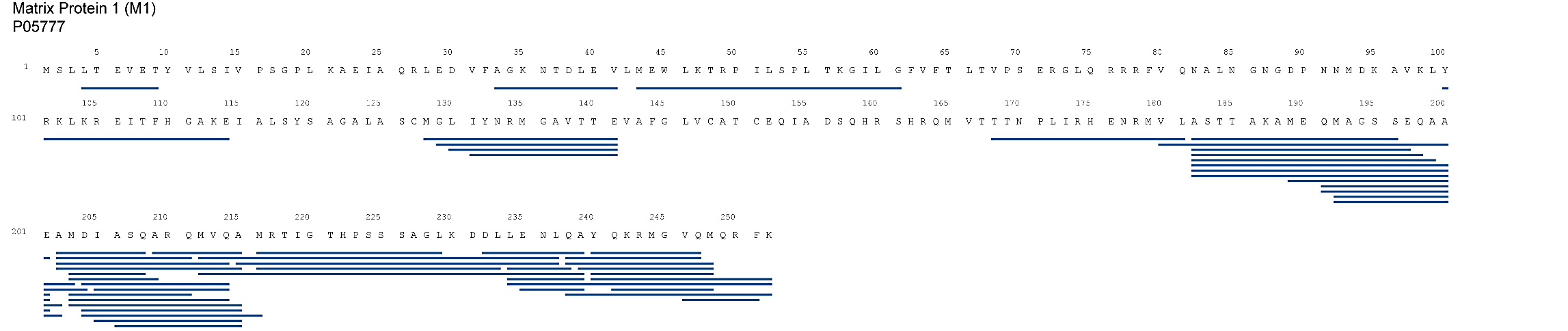

**Supplementary Figure S6 – HDX-MS peptide coverage map displaying unique peptides for influenza A virus M1.** Whole virus samples were acidified to pH 4, then neutralized back to pH 7 after 30 seconds. Samples were then incubated in D_2_O, quenched and processed for analysis by HDX-MS. Peptides aligned to published M1 protein sequence for strain A/WSN/33 (UniProt: P05777). Only unique peptides where HDX-MS data was attained for acid-treated samples are shown here. In these conditions, 68% of the M1 protein was covered by at least 1 unique peptide. Coverage of the N-terminal region was limited compared with the C-terminal, which is attributed to proximity of the N-terminal region to the viral envelope and potential protease inhibition. Coverage maps made using HDExaminer (Sierra Analytics).

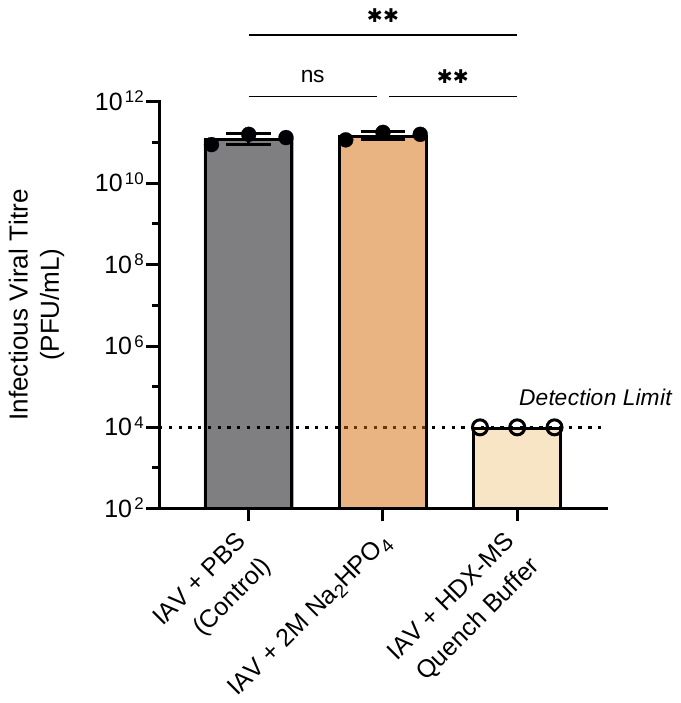

**Supplementary Figure S7** **– IAV infectious titer is not impacted by pH neutralization method.** Purified A/WSN/33 IAV stock (7.5 x 10^11^ PFU/mL) was mixed with 2M Na_2_HPO_4_ to mimic the pH neutralization process, in triplicate. Samples were vortexed to mix, then diluted 1/100 in PBSi for freezing, prior to titration by plaque assay. Control virus was mixed with an equivalent volume of PBS in the absence of Na_2_HPO_4_, then diluted in PBSi, frozen, and titrated in an identical manner. Virus samples were also mixed with ice-cold HDX-MS Quench buffer (150mM tris(2-carboxyethyl) phosphine-HCl, 2M urea, 0.1% formic acid final concentrations), vortexed briefly to mix, and incubated on ice for 2 min. Samples were then diluted 1/100 in PBSi for freezing prior to titration. Dotted line indicates detection limit of plaque assay, open symbols indicate data points below this limit. Data analyzed by One-Way ANOVA (ns, no significant difference, ** p < 0.01), n = 3 individual samples per treatment group.

**Supplementary Table S1** – Statistical results from Ordinary Two-Way ANOVA with Tukey’s Multiple Comparisons Test and a single pooled variance. Values here are related to Figure 2D, showing D_2_O incorporation into individual peptides from IAV protein Hemagglutinin (HA) with time. The summary of significance (Sig.) and adjusted P-values for each individual comparison are reported. Values indicate the statistical comparison of means from triplicate samples exposed to pH 7 or pH 4 (variable 1) for 2 different time periods (variable 2). Statistical comparisons are grouped according to the incubation time of samples in deuterium oxide (D_2_O, 3 different incubation times utilized for analysis of structural changes).

| **TWO-WAY ANOVA RESULTS** | **HA1 Peptide**  **51-58** | | **HA1 Peptide**  **68-79** | | | **HA1 Peptide**  **81-92** | | | **HA1 Peptide**  **89-95** | | | **HA1 Peptide**  **113-117** | | | **HA1 Peptide**  **317-328** | | **HA2 Peptide**  **521-532** | |
| --- | --- | --- | --- | --- | --- | --- | --- | --- | --- | --- | --- | --- | --- | --- | --- | --- | --- | --- |
| *with Tukey’s Multiple Comparisons Test* | *Sig.* | *P- Value* | | *Sig.* | *P-Value* | *Sig.* | *P- Value* | *Sig.* | | *P- Value* | *Sig.* | | *P- Value* | *Sig.* | | *P- Value* | *Sig.* | *P-Value* |
| **D_2_O - 30 sec** |  |  | |  |  |  |  |  | |  |  | |  |  | |  |  |  |
| pH 7 - 10 sec vs. pH 7 - 30 sec | ns | 0.6465 | | ns | 0.8535 | ns | 0.2907 | ns | | 0.9956 | ns | | >0.9999 | ns | | 0.485 | ns | 0.5725 |
| pH 7 - 10 sec vs. pH 4 - 10 sec | ns | 0.9862 | | * | 0.0115 | * | 0.0244 | ns | | 0.556 | **** | | <0.0001 | **** | | <0.0001 | **** | <0.0001 |
| pH 7 - 30 sec vs. pH 4 - 30 sec | ns | 0.0884 | | ns | 0.1024 | **** | <0.0001 | ns | | 0.5173 | **** | | <0.0001 | **** | | <0.0001 | **** | <0.0001 |
| pH 4 - 10 sec vs. pH 4 - 30 sec | ns | 0.7697 | | ns | 0.2798 | * | 0.0111 | ns | | 0.9909 | ns | | 0.9732 | ns | | 0.8309 | ns | 0.7823 |
| **D_2_O - 300 sec** |  |  | |  |  |  |  |  | |  |  | |  |  | |  |  |  |
| pH 7 - 10 sec vs. pH 7 - 30 sec | ns | 0.9696 | | ns | 0.2266 | ns | 0.9963 | ns | | 0.9021 | ns | | 0.9994 | ns | | 0.9997 | ns | 0.9919 |
| pH 7 - 10 sec vs. pH 4 - 10 sec | **** | <0.0001 | | ns | 0.096 | ns | 0.0947 | ns | | 0.0555 | **** | | <0.0001 | **** | | <0.0001 | **** | <0.0001 |
| pH 7 - 30 sec vs. pH 4 - 30 sec | **** | <0.0001 | | ns | 0.1763 | **** | <0.0001 | ** | | 0.0075 | **** | | <0.0001 | **** | | <0.0001 | **** | <0.0001 |
| pH 4 - 10 sec vs. pH 4 - 30 sec | ns | 0.5621 | | ns | 0.078 | * | 0.0121 | ns | | 0.9932 | ns | | 0.56 | ns | | 0.9988 | ns | 0.9998 |
| **D_2_O – 3,600 sec** |  |  | |  |  |  |  |  | |  |  | |  |  | |  |  |  |
| pH 7 - 10 sec vs. pH 7 - 30 sec | ns | 0.9994 | | ns | 0.2978 | ns | 0.9689 | ns | | 0.9873 | ns | | 0.549 | ns | | 0.9777 | ns | 0.9025 |
| pH 7 - 10 sec vs. pH 4 - 10 sec | **** | <0.0001 | | ns | 0.9997 | ns | 0.9999 | **** | | <0.0001 | **** | | <0.0001 | **** | | <0.0001 | **** | <0.0001 |
| pH 7 - 30 sec vs. pH 4 - 30 sec | **** | <0.0001 | | ns | 0.0791 | * | 0.0446 | **** | | <0.0001 | **** | | <0.0001 | **** | | <0.0001 | **** | <0.0001 |
| pH 4 - 10 sec vs. pH 4 - 30 sec | ns | 0.9766 | | ns | 0.8682 | ns | 0.1272 | ns | | 0.961 | ns | | 0.9965 | ns | | 0.9929 | ns | 0.9997 |

**Supplementary Table S2** – Statistical results from Ordinary Two-Way ANOVA with Tukey’s Multiple Comparisons Test and a single pooled variance. Values here are related to Figure 6C, showing D_2_O incorporation into individual peptides from IAV protein Matrix 1 (M1) with time. The summary of significance (Sig.) and adjusted P-values for each individual comparison are reported. Values indicate the statistical comparison of means from triplicate samples exposed to pH 7 or pH 4 (variable 1) for 2 different time periods (variable 2). Statistical comparisons are grouped according to the incubation time of samples in deuterium oxide (D_2_O, 3 different incubation times utilized for analysis of structural changes).

| **TWO-WAY ANOVA RESULTS** | **M1 Peptide**  **33-41** | | **M1 Peptide**  **100-114** | | | **M1 Peptide**  **128-141** | | | **M1 Peptide**  **131-141** | | | **M1 Peptide**  **168-181** | | | **M1 Peptide**  **182-197** | | | **M1 Peptide**  **216-229** | | | **M1 Peptide**  **234-239** | | | **M1 Peptide**  **240-252** | | |
| --- | --- | --- | --- | --- | --- | --- | --- | --- | --- | --- | --- | --- | --- | --- | --- | --- | --- | --- | --- | --- | --- | --- | --- | --- | --- | --- |
| *with Tukey’s Multiple Comparisons Test* | *Sig.* | *P-Value* | | *Sig.* | *P-Value* | | *Sig.* | *P-Value* | | *Sig.* | *P-Value* | | *Sig.* | *P-Value* | | *Sig.* | *P-Value* | | *Sig.* | *P-Value* | | *Sig.* | *P-Value* | | *Sig.* | *P-Value* |
| **D_2_O - 30 sec** |  |  | |  |  | |  |  | |  |  | |  |  | |  |  | |  |  | |  |  | |  |  |
| pH 7 - 10 sec vs. pH 7 - 30 sec | ns | 0.3004 | | ns | 0.8953 | | ns | 0.1325 | | ns | 0.4019 | | ns | 0.8556 | | ns | 0.6898 | | ns | 0.7543 | | ns | 0.6888 | | ns | 0.5155 |
| pH 7 - 10 sec vs. pH 4 - 10 sec | ns | 0.152 | | ns | 0.9877 | | * | 0.0395 | | ns | 0.0994 | | ns | 0.8438 | | ns | 0.5493 | | ns | 0.8449 | | *** | 0.0002 | | ns | 0.0521 |
| pH 7 - 30 sec vs. pH 4 - 30 sec | ns | 0.8994 | | ns | 0.1723 | | ** | 0.0095 | | * | 0.0299 | | ns | 0.2553 | | **** | <0.0001 | | **** | <0.0001 | | **** | <0.0001 | | **** | <0.0001 |
| pH 4 - 10 sec vs. pH 4 - 30 sec | ns | 0.9147 | | ns | 0.2863 | | * | 0.0371 | | ns | 0.1629 | | ns | 0.9921 | | **** | <0.0001 | | *** | 0.0001 | | **** | <0.0001 | | **** | <0.0001 |
| **D_2_O - 300 sec** |  |  | |  |  | |  |  | |  |  | |  |  | |  |  | |  |  | |  |  | |  |  |
| pH 7 - 10 sec vs. pH 7 - 30 sec | ns | 0.9995 | | ns | 0.7973 | | ns | 0.941 | | ns | 0.9974 | | ns | 0.9251 | | ns | >0.9999 | | ns | 0.4184 | | ns | >0.9999 | | ns | 0.8783 |
| pH 7 - 10 sec vs. pH 4 - 10 sec | ns | 0.9861 | | ns | 0.9355 | | ns | 0.7857 | | ns | 0.6237 | | ns | 0.3704 | | *** | 0.0005 | | * | 0.024 | | **** | <0.0001 | | ** | 0.0098 |
| pH 7 - 30 sec vs. pH 4 - 30 sec | ns | 0.9204 | | ns | 0.9712 | | **** | <0.0001 | | *** | 0.0004 | | ns | 0.1009 | | **** | <0.0001 | | ** | 0.0033 | | **** | <0.0001 | | **** | <0.0001 |
| pH 4 - 10 sec vs. pH 4 - 30 sec | ns | 0.9773 | | ns | 0.8925 | | ** | 0.0018 | | * | 0.0112 | | ns | 0.5214 | | **** | <0.0001 | | * | 0.0259 | | **** | <0.0001 | | ** | 0.0096 |
| **D_2_O – 3,600 sec** |  |  | |  |  | |  |  | |  |  | |  |  | |  |  | |  |  | |  |  | |  |  |
| pH 7 - 10 sec vs. pH 7 - 30 sec | ns | 0.75 | | ns | 0.0728 | | ns | 0.8682 | | ns | 0.5594 | | ns | >0.9999 | | ns | 0.9931 | | ns | 0.8614 | | ns | 0.9989 | | ns | 0.9997 |
| pH 7 - 10 sec vs. pH 4 - 10 sec | ns | 0.8569 | | ns | 0.7913 | | ** | 0.0061 | | ** | 0.0015 | | ** | 0.0037 | | **** | <0.0001 | | ns | 0.2344 | | * | 0.0106 | | ns | 0.5318 |
| pH 7 - 30 sec vs. pH 4 - 30 sec | ns | 0.2391 | | ns | 0.8292 | | *** | 0.0004 | | *** | 0.0005 | | **** | <0.0001 | | **** | <0.0001 | | *** | 0.0003 | | **** | <0.0001 | | ** | 0.0078 |
| pH 4 - 10 sec vs. pH 4 - 30 sec | ns | 0.4029 | | ** | 0.0098 | | ns | 0.2498 | | ns | 0.3105 | | *** | 0.0002 | | **** | <0.0001 | | ns | 0.1374 | | **** | <0.0001 | | ns | 0.1774 |

**Supplementary Tables S3, S4, S5** – External Excel tables listing HDX-MS raw data including the list of peptide analyzed, deuteration levels and differences in deuteration levels for each timepoint. One table is included for each protein studied (HA; NP; M1).
